# Biologically informed deep learning to infer gene program activity in single cells

**DOI:** 10.1101/2022.02.05.479217

**Authors:** Mohammad Lotfollahi, Sergei Rybakov, Karin Hrovatin, Soroor Hediyeh-zadeh, Carlos Talavera-López, Alexander V Misharin, Fabian J. Theis

**Author notes:** equal contribution (Mohammad Lotfollahi, Sergei Rybakov).

## Abstract

The increasing availability of large-scale single-cell datasets has enabled the detailed description of cell states across multiple biological conditions and perturbations. In parallel, recent advances in unsupervised machine learning, particularly in transfer learning, have enabled fast and scalable mapping of these new single-cell datasets onto reference atlases. The resulting large-scale machine learning models however often have millions of parameters, rendering interpretation of the newly mapped datasets challenging. Here, we propose expiMap, a deep learning model that enables interpretable reference mapping using biologically understandable entities, such as curated sets of genes and gene programs. The key concept is the substitution of the uninterpretable nodes in an autoencoder’s bottleneck by labeled nodes mapping to interpretable lists of genes, such as gene ontologies, biological pathways, or curated gene sets, for which activities are learned as constraints during reconstruction. This is enabled by the incorporation of predefined gene programs into the reference model, and at the same time allowing the model to learn *de novo* new programs and refine existing programs during reference mapping. We show that the model retains similar integration performance as existing methods while providing a biologically interpretable framework for understanding cellular behavior. We demonstrate the capabilities of expiMap by applying it to 15 datasets encompassing five different tissues and species. The interpretable nature of the mapping revealed unreported associations between interferon signaling via the RIG-I/MDA5 and GPCRs pathways, with differential behavior in CD8^+^ T cells and CD14^+^ monocytes in severe COVID-19, as well as the role of annexins in the cellular communications between lymphoid and myeloid compartments for explaining patient response to the applied drugs. Finally, expiMap enabled the direct comparison of a diverse set of pancreatic beta cells from multiple studies where we observed a strong, previously unreported correlation between the unfolded protein response and asparagine N-linked glycosylation. Altogether, expiMap enables the interpretable mapping of single cell transcriptome data sets across cohorts, disease states and other perturbations.

## Introduction

The progress and development of experimental technologies^1–4^ and computational tools^5–9^ for single-cell genomics have enabled the construction of atlases with millions of cells serving as high-resolution coordinate systems^10^ for biological and therapeutic discoveries^11–14^. However, leveraging existing atlases poses a computational challenge known as reference mapping. Reference mapping allows the rapid integration of newly generated datasets, denoted as a query, hereby facilitating their analysis and interpretation. The transfer of knowledge from the reference to the query allows the rapid annotation of the query data^7^, imputation of missing modalities in the query^8,15^, and the discovery of novel populations, such as disease states unseen in the reference dataset^8,15^.

Single-cell reference mapping is growing in popularity^16^, with approaches such as Seurat’s supervised principal component analysis^8^, our single-cell architecture surgery (scArches)^15^, and an extension of Harmony^17^ to map query datasets by minimal modification of the reference atlas^18^. Existing reference mapping methods embed new query data into a reference latent space by removing technical differences such as batch effects between the reference and the query without access to reference data. However, the implicitly used latent dimensions for joint data representation are not directly interpretable. An important trend in machine learning is the development of interpretable models, e.g., by the addition of statistical assumptions to learned latent spaces or the inclusion of prior information from validated mechanisms or other data^19^. As the former disentanglement approaches have not yielded sufficiently useful latent spaces in our context^20–22^, we here hypothesize that using prior information instead may help identifiability. In particular, we aim to leverage known or newly learned gene programs (GPs) to contextualize query data by answering various questions, including “which GPs are disturbed in a disease query data compared to the healthy reference?” and “which biological programs explain a novel population in the query?” By thus making reference mapping interpretable, it can move beyond mere data alignment between query and reference and be used for further interpretation of query data for example in the case of disease perturbation versus a healthy atlas. Currently, the standard approach for the identification of biological programs in query cells compared with a reference atlas is to test for differentially expressed genes and downstream gene set enrichment. However, differential expression on an atlas consisting of cells from an arbitrary number of studies with variable degrees of biological and technical heterogeneity represents a challenge for the statistical analysis. The currently accepted best practices^23,24^ suggest that differential expression should be performed on non-integrated expression data and not on the corrected expression values after integration; hence, statistical models should account for complex experimental designs and adjust for unwanted variation, such as batch effects. Moreover, modeling constraints, such as inestimable coefficients and less than full rank models, can hamper the proper assessment of differentially expressed genes. In contrast, the simpler non-parametric statistical tests potentially capture both biologically relevant and irrelevant genes, which may compromise the accuracy of enriched gene set terms.

Collectively, it may therefore be useful to have interpretable embeddings directly associated with validatable GPs in the context of atlas-wide comparisons to capture the relevant biological signals while accounting for nonlinear batch effects. This end-to-end approach is common in deep learning and has been shown to outperform classical approaches that use sequential regularization and analysis^19^. Interpretable reference mapping requires the incorporation of domain knowledge^19^, such as curated GPs, into the representation learning model to guide interpretation and exploration. The inclusion of domain knowledge to design “domain-informed” deep learning architectures has been shown to improve the performance on challenging prediction tasks, from tumor type^25^ to protein structure^26^. However, the methods to include such knowledge, while accounting for the incompleteness of knowledge bases and remaining flexible to discover novel insights, rather than being locked into prior-based feature design, remain difficult.

To address these challenges, we propose to build a machine learning system that exploits the knowledge of the underlying biological phenomenon for single-cell representation learning (as outlined more generally in the idea of “differential programs”^19^ recently). We construct an *“explainable programmable mapper” (expiMap)* as an interpretable conditional variational autoencoder^7,27,28^ (CVAE) that allows the incorporation of domain knowledge by performing “architecture programming”, i.e., constraining the network architecture to ensure that each latent dimension captures the variability of known GPs. We apply an attention-like mechanism^29^ to select the relevant GPs for each reference dataset. This helps with the prioritization of significant gene sets, but also allows the inclusion of genes that were not originally included in annotated GPs, thereby addressing the incomplete nature of the knowledge database. To identify new variations unique to the query data, such as diseases effects, we identify *de novo* GPs in addition to the known GPs in knowledge bases, by learning disentangled latent representations. Through multiple examples, we demonstrate that expiMap provides an interpretable reference mapping framework to answer program-level queries. The framework can be used for the discovery of novel insights into the biological processes in normal and disease states directly when mapping to the atlas, while maintaining comparable integration performance to existing data integration methods.

## Results

### Interpretable single-cell reference mapping using expiMap

Linear methods, such as PCA^30,31^ or matrix factorization^32,33^, learn a representation of the data where each dimension of the latent space can be explained using a weighted combination of the input, such as gene expression. This interpretability comes at the cost of the limited capacity (e.g., only capturing linear relationships) of the model to fit the data. In contrast, nonlinear methods using deep neural networks^34,35^ come with a larger capacity at the expense of reduced model interpretability.

Here, we aim to design a system with the ability to provide biologically interpretable answers to queries of an integrated representation of multiple (denoted by N) reference single-cell datasets and custom “gene programs.” These can be gene lists from existing curated databases^36,37^, lists extracted from literature^38^ or individually curated gene sets (**Fig. 1a**). This knowledge is transformed into a binary GP matrix, in which each row is a gene program, and each column denotes the membership of a gene in that program (see **Methods** and **Fig. 1b**).

**Fig. 1.**
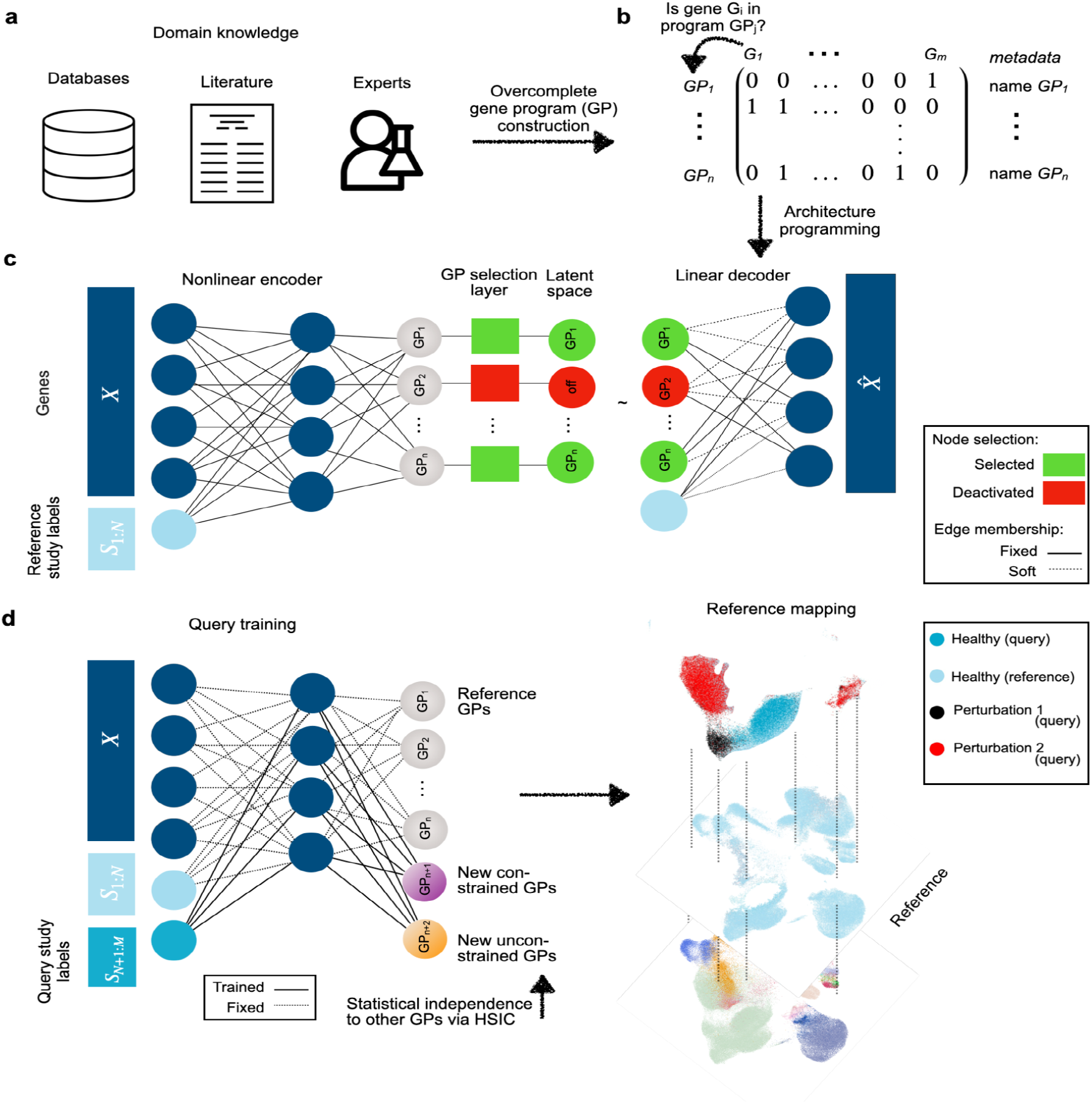
Biologically informed reference mapping using expiMap. **(a)** Domain knowledge from databases, articles, and expert knowledge is used to construct a binary matrix of gene programs (GPs) shown in **(b). (c)** The model is trained on reference data, receiving gene expression and study label for each cell to encode a set of latent variables representing GPs. The GPs are pruned and enriched by the model using a group lasso and gene-level sparity regularization, respectively and fed into a linear decoder. The GP matrix is then used to program the neural network architecture by wiring the model parameters of the decoder to learn a specific GP for each latent dimension. **(d)** Upon mapping query data, the reference model is expanded and fine-tuned using architecture surgery, whereas new learnable latent GPs are added and trained with the query data. The decoder architecture equals **(c)** with the difference that only highlighted weights of newly added GPs are trainable in the encoder and decoder. To make sure these newly learnt unconstrained GPs do not overlap with reference GPs, we employ statistical independence constraints. HSIC: Hilbert–Schmidt independence criterion.

We then propose *architecture programming* i.e. wiring the weights of the network using the GP matrix such that each latent variable contributes to the reconstruction of a set of genes defined by the GP. The model receives a gene expression matrix from *N* different single-cell studies (*X*) and an additional vector for corresponding one-hot encoded study labels (*S*_1:*N*_) for each cell, for example the experimental laboratories or sequencing technologies **(Fig. 1c).** The adopted variational autoencoder architecture^7,34^ leverages a nonlinear encoder for flexibility and a linear decoder for interpretability. The latent space dimension chosen is equal to the number of GPs, and the weights from each latent dimension (i.e., latent GP) to output are programmed according to the GP matrix so that a latent GP can only contribute to the reconstruction of genes in a particular GP (denoted as “fixed membership” in **Fig. 1c**). Since annotated GPs are often incomplete, we allow the inclusion of other genes in each GP by applying L1 sparsity regularization to genes not initially labeled to belong to that GP (denoted as “soft membership” in **Fig. 1c**). This enables the model to leverage the sparse selection of other genes, which helps in the reconstruction and therefore accounts for incomplete domain knowledge, to refine ontologies and pave the way toward a data-driven alternative means to learn gene programs (see later results).

However, the number of GPs may be very large, potentially redundant, and not all of them are relevant for every atlas. To select only informative GPs, an attention-like mechanism is implemented with a group lasso regularization layer in latent space (see **Methods**), which deactivates GPs that are redundant or do not contribute to the reconstruction error of the model. The model is trained in an end-to-end manner and can thus be used to construct reference atlases with interpretable embedding dimensions, which we can leverage to analyze integrated datasets.

Based on this pretrained, interpretable reference model, we now propose to employ transfer learning, as outlined in architectural surgery^15^ (see **Methods**), to map new datasets into the reference. We modify the strategy of fine-tuning conditional weights in scArches allowing the model to learn new GPs that are not included in the reference model. This is achieved by the addition of new latent space dimensions, i.e., nodes with trainable weights in the bottleneck layer of the model (**Fig. 1d,**see **methods**), while keeping the rest frozen. We implement two ways of learning these new GPs: either by learning GPs confined to predefined genes (denoted as “new constrained” in **Fig. 1d**) that were not present or those that have been deactivated in the reference model. In addition, the model may also learn *de novo* GPs as realized by an L1-regularized gene prior, to capture new variations in the query data without predefined gene sets (denoted as “new unconstrained” in **Fig. 1d**).The limited learning capacity of the model at the reference mapping stage, due to frozen weighting, enforces an information bottleneck (i.e., a reduced capacity to learn and store information), encouraging the new nodes to learn important and potentially disentangled^39^ sources of variations in the query data. We further employ the Hilbert–Schmidt Independence Criterion (HSIC)^21,40^, a kernel-based measure of latent variable independence^40^, to enforce independence between old and new unconstrained GPs learned during query optimization (**Fig. 1d**).

The probabilistic representation learned by expiMap as a Bayesian model allows the performance of hypothesis testing on the integrated latent space of the query and the reference accounting for technical factors (see **methods**).The hypothesis testing is performed at the GP level, enabling the identification of differential GPs between two groups of cells by sampling from the group’s posterior distribution of the latent variables. The ratio between two hypothesis probabilities is reported by the Bayes factor. Later, we demonstrate how this ability helps to identify GPs associated with perturbation in the query data compared with the healthy reference.

Collectively, through expiMap, we propose an approach to learn interpretable, domain-aware representations of single-cell datasets for the integrative analysis of reference and query data. Further, we propose a modified version of architecture surgery that goes beyond predefined domain knowledge while retaining interpretability. This allows the contextualization of the query data with the reference data within a specific GP to provide answers to the user’s biological questions.

### expiMap parses single-cell transcriptional response to interferon-beta

One of the ultimate goals in building large, single-cell atlases is the study of the effect of perturbations (e.g., disease) and contextualizing it within a given reference. To demonstrate the applicability of our model in this scenario, we constructed a human immune cell atlas from four different studies of bone marrow^41^ and peripheral blood mononuclear cells (PBMCs)^42–44^. We then mapped a query PBMC dataset of samples from eight patients diagnosed with systemic lupus erythematosus whose cells were either untreated (control) or treated with interferon (IFN-β), a potent cytokine inducing a strong transcriptional response in immune cells^45^. Successful mapping should align untreated cells to matching cell types in the healthy reference while preserving the strong effect of IFN-β. The expiMap model trained with GPs extracted from the Reactome^36,37^ pathway knowledgebase successfully mapped the query untreated cells to the healthy reference while forming clusters indicative of the IFN-β-treated cells (**Fig. 2a**).

**Fig. 2.**
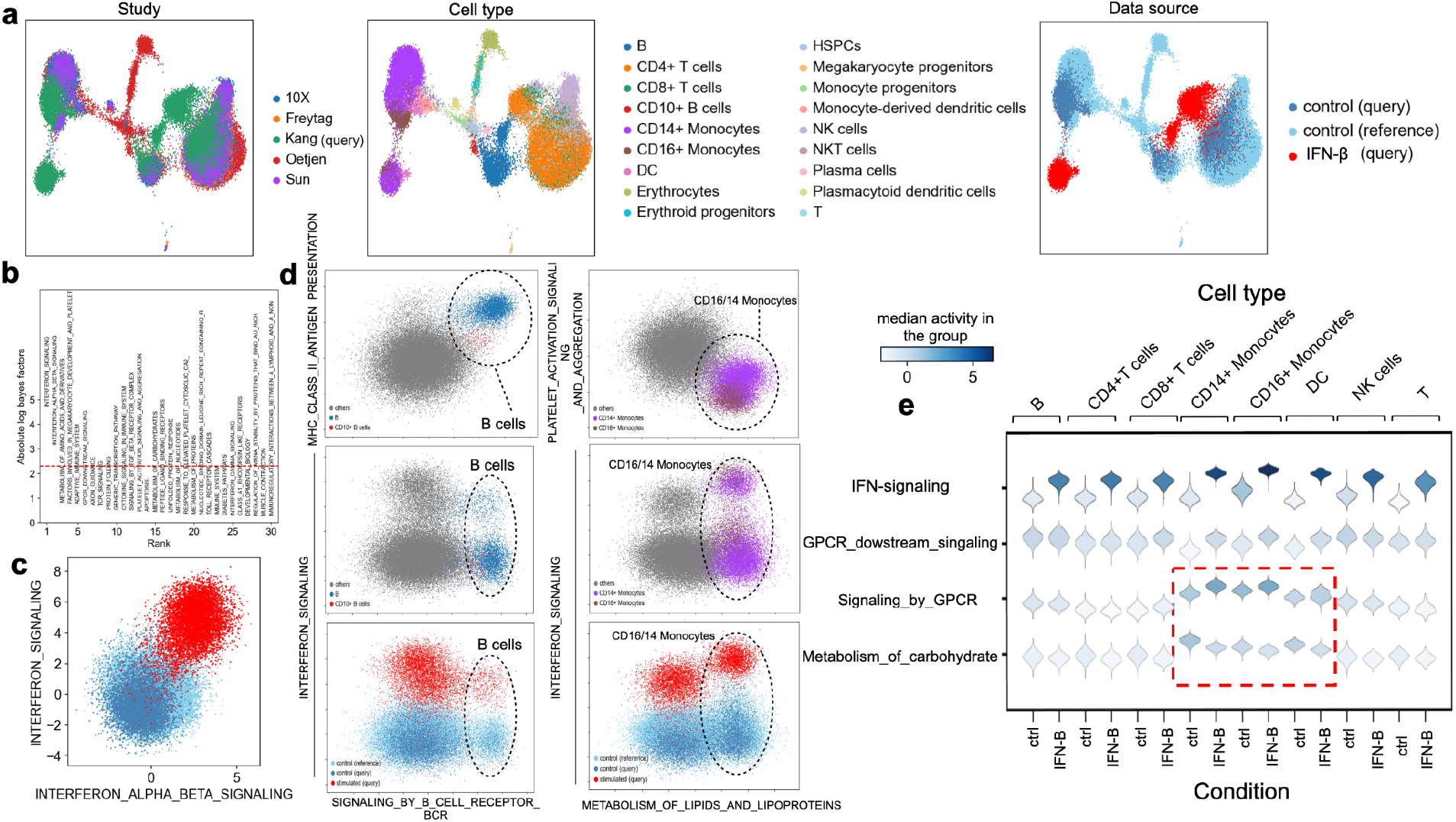
ExpiMap resolves gene programs after IFN-β perturbation. **(a)** UMAP representation of the query control and IFN-β stimulated cells from eight patients (n = 13,576 cells) mapped onto a healthy immune reference from four different studies (n = 32,484 cells) using expiMap. Colors demonstrate study (left), harmonized cell type (middle), and data source (right). **(b)** Differential GP analysis results between query IFN-β and control cells from both the query and reference. The x-axis shows the ranking of GPs; the y-axis denotes the significance (absolute log-Bayes factor) of each GP. **(c)** Visualization of both the reference and query data in the context of the top two most significant expiMap latent GPs in **(b)**. Each dot shows the latent GP score of each cell. **(d)** Visualization of the query and reference in various GPs, delineating cell types or perturbation states for B cells and CD14^+^/16^+^ monocytes. **(e)** The activity of the most differentially active GP terms in CD14^+^ monocytes after IFN-β stimulation. Each violin plot demonstrates the distribution of latent GP values across different cell types. The dashed square highlights GPs characterizing the myeloid-specific response to IFN-β.

By testing between IFN-β and control conditions, we identified the top differential GPs, matching to previously reported GPs^46,47^ including interferon-related pathways (**Fig. 2b**),which also separates the control reference and query cells from stimulated query cells (**Fig. 2c**).Following up with a cell type-specific analysis, we identified differential GPs across cell types (i.e., one vs all) or cell type-specific IFN-β effects (i.e., IFN-β vs control within a cell type). In particular, we detected a group of population-specific GPs that separated one cell type from the rest (**Fig. 2d, first row**).The population-specific GPs can be used together with perturbation-associated GPs (i.e., obtained from IFN-β query cells vs control cells in both query and reference for that cell type) to resolve the heterogeneity of cell state for that cell type (**Fig. 2d, second row;**see **Supplementary Figs. 1–3** for all cell types). We found that the general interferon GPs (e.g., interferon signaling) are always induced in all cell types (see **Fig. 2f** and **Supplementary Fig. 3**), whereas some GPs (e.g., GPCR-related programs), including genes from the CXC chemokine family (e.g., CXCL10), are only present in the myeloid lineage (see highlighted GPs in **Fig. 2d;** see **Supplementary Fig. 3** for all extended figures). Additionally, we detected carbohydrate metabolism activity in CD14^+^ and CD16^+^ monocytes and DCs, and active amino acid metabolism in CD14^+^ monocytes after IFN-β stimulation (**Supplementary Fig. 3**).This is in agreement with previous observations in cancer and viral infection showing that amino acid, lipid, and carbohydrate metabolic pathways contribute to the immune response^48,49^. Specifically, it is known that IFN-β engages with the amino acid metabolic pathway to produce polyamines and clear viral infections^50^, but a direct link to myeloid cells, as revealed by expiMap, has not been described elsewhere.

Differential expression analysis on atlases is challenging due to the complex experimental designs and probable presence of nonlinear batch effects that cannot be modeled by linear approaches. Gene Set Enrichment Analysis (GSEA) is a classical approach for inferring the activity of gene programs and involves the sequential pipeline of differential expression analysis and gene set enrichment test. To evaluate robustness of expiMap’s integrated GP test, we hence compared it to the classical GSEA via limma-fry^100,101^ (see **Supplementary note 1** and **Supplementary Fig. 4).**In our comparisons, we observed that unlike conventional gene set testing, which tends to detect general, non-specific terms, expiMap was able to identify specialized GPs. For example, in the B cell population of both IFN-β-treated and control cells, expiMap detected B cell receptor signaling and antigen presentation activity, which are more descriptive of B cell biology than the general terms such as “adaptive immune response” or “immune response” that were found to be enriched in these cells by limma-fry (see **Supplementary Fig. 4c**).We postulate that the increased variability in gene expression measurements hinders the detection of specialized biological signals by standard gene set testing on cell atlases. This indicates that expiMap can extract biologically relevant GPs from a single-cell atlas consisting of many datasets while accounting for technical variations such as batch effects, which may not always be feasible with existing pipelines, owing to the presence of nonlinear batch effects.

### Domain awareness improves the performance of simpler models

As a means to benchmark the performance of expiMap’s reference mapping component, we compared it to scArches + scVI^7^, Seurat v4^8^, and Symphony^17^. Although expiMap and scVI both leverage scArches for reference mapping, scVI did not mix the untreated monocytes from the query data with healthy monocytes in the reference (dotted circle in **Fig. 3a**; see **Supplementary Fig. 5** for mixing of studies), whereas expiMap successfully integrated them into the healthy reference (0.68 vs 0.47 average batch correction scores; see further for description of the metrics) while preserving the effect of IFN-β treatment in cells that should not be integrated with the rest. We attribute this to the explicit incorporation of the IFN-β related GPs in the expiMap model, which helps to differentiate the perturbed and control states while resolving the transcriptional similarities between control cells, hence leading to a better mixing of control states. We investigated this by removing the top five GPs obtained from the IFN-β vs control comparison (see **Fig. 2b**) and retraining the model. We observed that this led to the incorrect mixing of control and stimulated cells with the reference (see **Supplementary Fig. 6**).In this example, both scArches + scVI and expiMap had better performance than Seurat V4 and Symphony for integrating control query cells into control cells from the reference (**Fig. 3b**).We also quantitatively evaluated the integration of query control cells into the healthy reference using nine different metrics of biological preservation and mixing^51^.

**Fig. 3.**
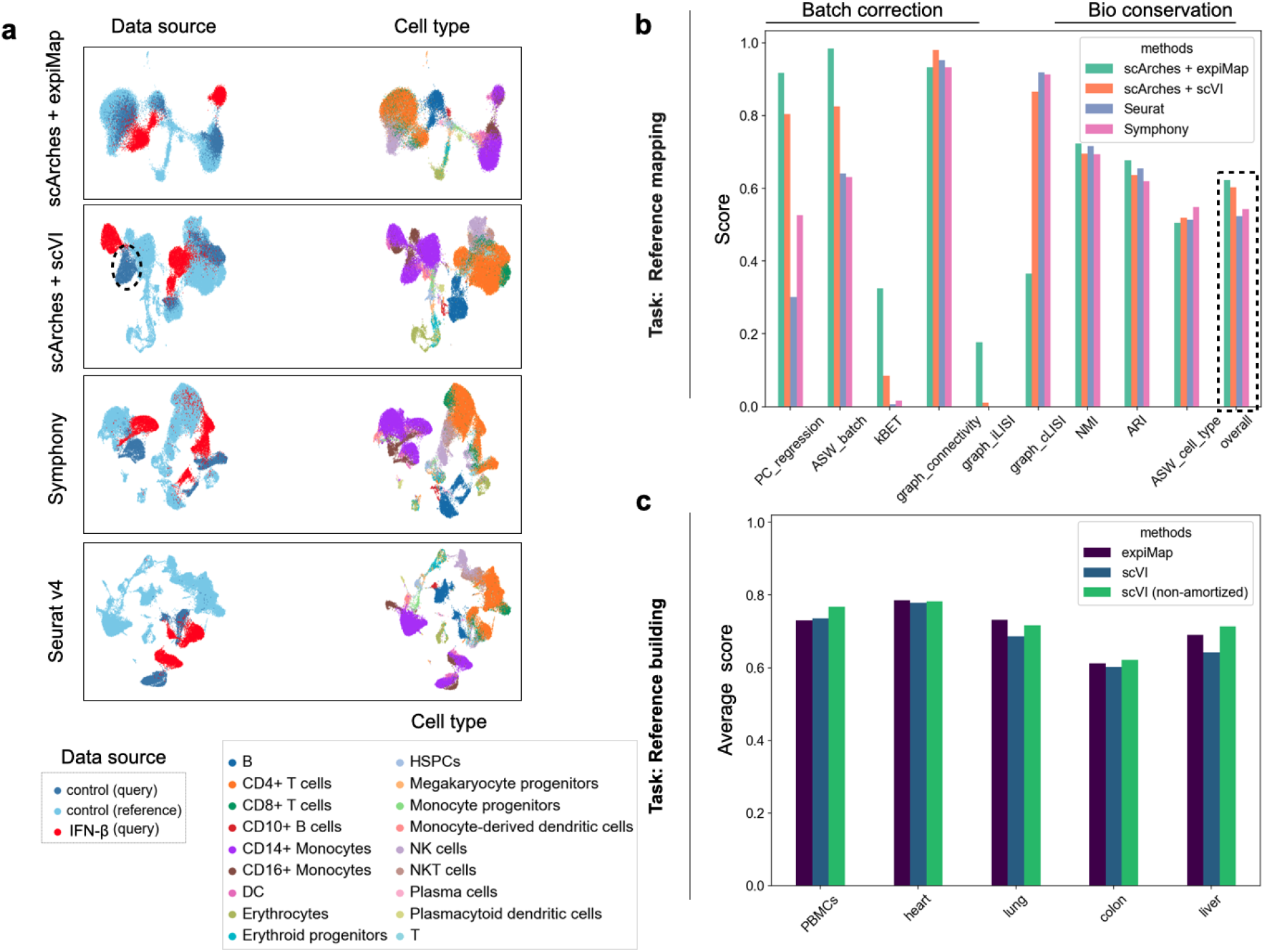
Domain awareness improves performance in downstream tasks. **(a)** UMAP representation of integrated healthy immune reference with query interferon IFN-β data from eight patients for expiMap and existing reference mapping methods. Colors denote the data source and cell type. The dotted circle highlights query control monocytes that scArches + scVI failed to integrate into the control reference. **(b)** Comparison of integration accuracy for mapping <u>control</u> query cells (excluding IFN-β cells) onto healthy atlases across different models. The metrics measure batch correction and bioconservation. The dotted line is the overall score calculated based on the mean of all metrics. **(c)** expiMap retains the expressiveness of an unconstrained reference model, as shown by the comparison of reference building performance through benchmarking in five different tissues, including PBMCs (n = 161,764, n_batches = 8), heart (n = 18,641, n_batches = 4), lung (n = 65,662, n_batches = 19), colon (n = 34,772, n_batches = 12), and liver (n = 113,063, n_batches = 14) across three different methods. The y-axis is the average score of the nine metrics detailed in **(b).** PC regression, principal component regression. kNN, k-nearest neighbor. ASW, average silhouette. ARI, adjusted Rand index. NMI, normalized mutual information.

The improvements achieved by reference mapping with expiMap compared with scVI as one of the top performers in atlas-level integration benchmarks^51^ motivated us to investigate this performance gain further. While both scVI and expiMap are variational autoencoders, scVI implements a nonlinear encoder and decoder whereas expiMap uses a linear and lower capacity decoder to enable interpretability. We hypothesized that the additional domain knowledge in expiMap combined with a linear decoder helps to find better posterior distributions in the encoder by improving a well-known problem called the “amortization gap“^52^, which is defined as the difference attributable to amortizing variational parameters over all the training data compared with the estimations for individual training samples. This leads to suboptimal variational approximation by the encoder in latent variable variational models^52^. This problem concerns the suboptimal posterior distribution learned. Thus, expiMap would find a richer representation within the family of all possible solutions compared with the less optimal representation found by scVI among more complex families of solutions enabled by additional non-linearities in the model. To test this, we modified the scVI encoder to employ a non-amortized formulation (see **Methods**), in which the parameters of the variational distribution are optimized for each cell individually. We trained expiMap, scVI, and non-amortized scVI to construct references with multiple atlases across five tissues obtained from Sfaira^53^. We observed that non-amortized versions of scVI consistently achieved superior or equal performance in data integration compared with expiMap, while remaining better than the default (amortized) scVI, which corroborated our hypothesis (**Fig. 3c**).We further performed similar benchmarking to evaluate our model against linear-decoded variational autoencoder (LDVAE)^54^, a variation of scVI with a linear decoder. We found that LDVAE had similar performance to amortized scVI, yet poorer performance than expiMap and non-amortized scVI, demonstrating the importance of including domain knowledge in expiMap (see **Supplementary Fig. 7).**

Overall, our results confirm that domain awareness enables expiMap to achieve state-of-the-art results for both reference mapping and reference construction. This is aligned with recent results^2,19^ demonstrating the improved performance of deep learning-based models by integrating domain knowledge into modeling.

### Learning novel interpretable programs to go beyond existing domain knowledge

Leveraging domain knowledge is crucial for the rapid and interpretable analysis of new query datasets within the context of a reference atlas. However, domain knowledge is not always comprehensive, complete, and up-to-date for a novel phenomenon (e.g., a new disease such as COVID-19). Thus, the ability to learn new GPs to analyze query data containing new variations, such as new states or cell populations, is pivotal. As explained in the first results section, we address this by allowing expiMap to learn novel GPs associated with the query data that exist in the knowledge base but are not detected previously in the reference model, as well as *de novo* programs that are not described in the knowledge base (see **methods**).To evaluate the success of this strategy, we sought to remove GPs and cells containing information about interferon signaling and B cells during reference training and assess if the model could *de novo* learn GPs of that type if the query data contain B cells and IFN-β treated cells. To this end, we removed the general IFN-related GPs, including IFN, IFN-αβ (and GPs containing a superset of those), and cytokine signaling in the immune system, from Reactome. We also removed B cells in the reference and the top two B cell GPs containing information about B cell receptor signaling and antigen presentation, as shown in **Fig. 2d.** Next, we trained the healthy reference PBMC model, as before, with the same studies as **Fig. 2a,**in which the model did not see gene programs related to IFN pathway activity, B cells, and their GPs in reference training. Further, we added a set of new nodes along with trainable weights at the query training stage; one of them was set with fixed gene membership to learn B cell receptor signaling GP and the other three were flexible and able to learn other variations in the data. In practice, we suggest initializing ten (as default) newly initialized unconstrained nodes for more complicated query datasets as redundant nodes will be switched off (all decoder weights set to zero) by L1 regularization. Ideally, we would like the model to learn GPs containing information about new variations in the query. We examined the distribution of the latent space values across different cell types (**Fig. 4a**).The node that was constrained with the B cell GP indeed learned the variations specific to B cells (**Fig. 4a,**first row). The B cell node had 84 active genes, of which 66 genes are from the B cell receptor signaling GP (**Fig. 4b**).While expiMap learned the predefined GP, it also added nine B cell markers (see **Supplementary Table 1** for the full gene list) obtained using differential testing (Wilcoxon rank-sum test in scanpy^55^) owing to the soft membership features in the model that were not initially in the predefined GP, demonstrating the ability of the model to incorporate extra information and enrich incomplete domain knowledge (**Fig. 4b**).Further, by looking at distribution plots, one of the newly learned nodes after in query training displayed a different distribution for myeloid cells/lineage (denoted as Node 1 **Fig. 4a**), while other cells had uniformly similar values. Another node (Node 2 **Fig. 4a**) had a bimodal distribution across all cell types, which potentially suggests that the variation between control and IFN-β stimulated cells are captured. Finally, the last node (Node 3 **Fig. 4a**) was deactivated for the most part of the training, capturing no clear variation (**Fig. 4b** last row and see **Supplementary Fig. 8**).We then sought to uncover the variations in the *de novo* learned nodes by comparing the top 50 genes influencing that node (see **methods** for ranking scheme) with GPs with maximum number of overlapping genes and those from differentially expressed genes. We found that Node 1 and Node 2 learned variations related to myeloids and IFN-β (**Fig. 4c**).Specifically, Node 1 is a new GP with minimal overlap with the top two previously identified programs (**Supplementary Fig. 9**).This newly learned program also had a maximum gene overlap of 24% with the top 50 genes influencing the GP with other existing GPs in Reactome (GPs that had maximum gene overlap are shown on the first row in **Fig. 4c**).This demonstrates that the model learned a new program distinguishing myeloid cells from other cell types. Additionally, Node 2 also captured the program describing the interferon response only observed in the query data. When plotted against each other, we observe the separation of B cells and myeloid cells **(Fig. 4d),**IFN-β treated cells and B cells (**Fig. 4 e–f**).As we demonstrated for B cells, expiMap can enrich predefined and potentially incomplete GPs. We further examined this feature by incrementally removing the most influential genes from general interferon signaling GP-trained, IFN-β-treated cells, and control cells from the data of Kang et al., while also monotonically increasing the L1 sparsity. Lower L1 values encourage the model to add more genes to the predefined GP compared with higher L1 values, therefore restricting the model to the predefined features. We observe that the model robustly recovers deleted genes with different ranges of L1 values (see **Supplementary Fig. 10a**); however, for lower L1 values, the model can also incorrectly add genes that may not be in the original program (see **Supplementary Fig. 10b**).

**Fig. 4.**
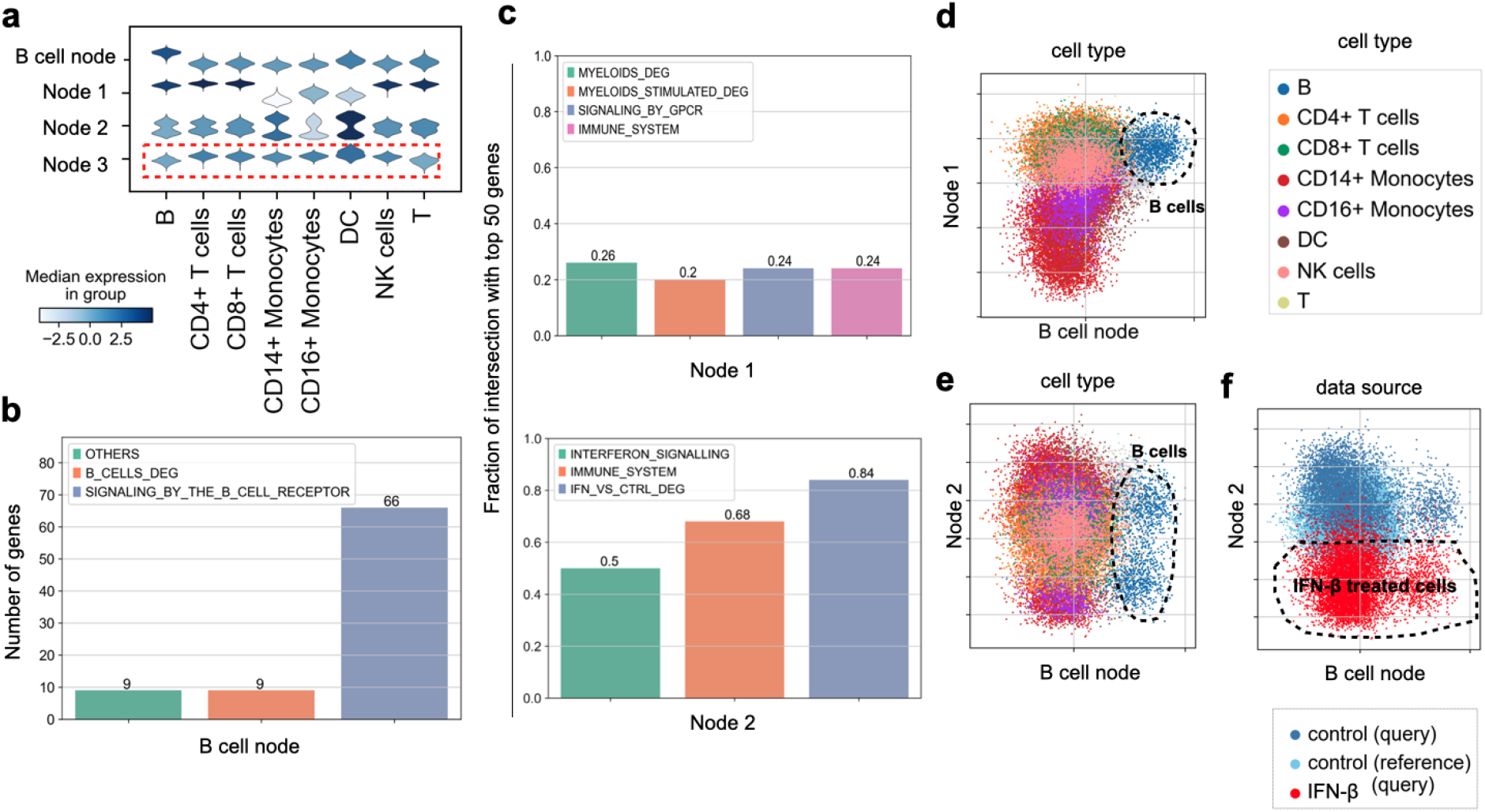
Learning new gene programs from query data. **(a)** Distribution of single-cell latent representation values across newly learned GPs across different query data cell types for query IFN-β treated cells and control cells. The highlighted row denotes a deactivated node capturing no strong discriminative variation between cell type or conditions. **(b)** Comparison of overlap of the most influential genes dominating the variance in newly learned constrained B cell nodes **(b)** and unconstrained nodes **(c)** with genes in existing related GPs and top genes obtained from the differential testing analysis. The terms “Myeloid_DEG” and “B_DEG” refer to genes obtained from one vs all Wilcoxon rank-sum tests in the query control cells for each population, respectively. The myeloid population is a collection of CD14^+^ monocytes, CD16^+^ monocytes, and DC populations. “INF_VS_CTRL_DEG” denotes differentially expressed genes comparing IFN-β treated and control cells. The existing GPs for **(c)**are those with maximal overlap with at least 12 genes with newly learned GPs. **(d–f)** Visualization of newly learned GPs discriminating specific cell types and states from the rest, such as B cells and myeloids with the effect of interferon removed **(d)** or B cells with the effect of interferon preserved **(e, f)**.

Overall, we demonstrated that expiMap can learn predefined GPs not in the reference GP matrix for populations only present in the query data during query training while having the ability to enrich the predefined GPs with new genes not in the program. In addition, we demonstrated that expiMap is not restricted to predefined GPs and can learn *de novo* GPs without any user supervision or prior knowledge. The *de novo* GPs are disentangled from previous GPs and can capture novel variations in the query data.

### Interpreting the immune response of patients with COVID-19 treated with an immunosuppressant

To demonstrate medical use of interpretable atlas querying, we aimed to determine the transcriptional programs of the cellular response to infection during COVID-19 and examine how they are affected by immunosuppressive interventions. We leveraged the integrated immune PBMC atlas to map IFN-β dataset (as in **Fig. 2a**) and a new dataset from two patients (P1 and P2) at different COVID stages (severe disease and during the remission process: D1, severe COVID-19 on day 1; D5 and D7, remission on days 5 and 7, respectively). Both patients were treated with tocilizumab, an immunosuppressive drug targeting the interleukin-6 (IL-6) receptor^56^. The integrated dataset (**Fig. 5b, c**) was reannotated using canonical markers identifying 20 cell states from the myeloid and lymphoid compartments, including rare populations such as megakaryocytes and erythroid progenitors, as well as a population of *CD10^+^* B cells (**Fig. 5c**).From the integrated embedding produced by expiMap, we could observe that some cellular states are associated with disease severity, which may be related to differences in the cellular response to tocilizumab.

**Fig. 5.**
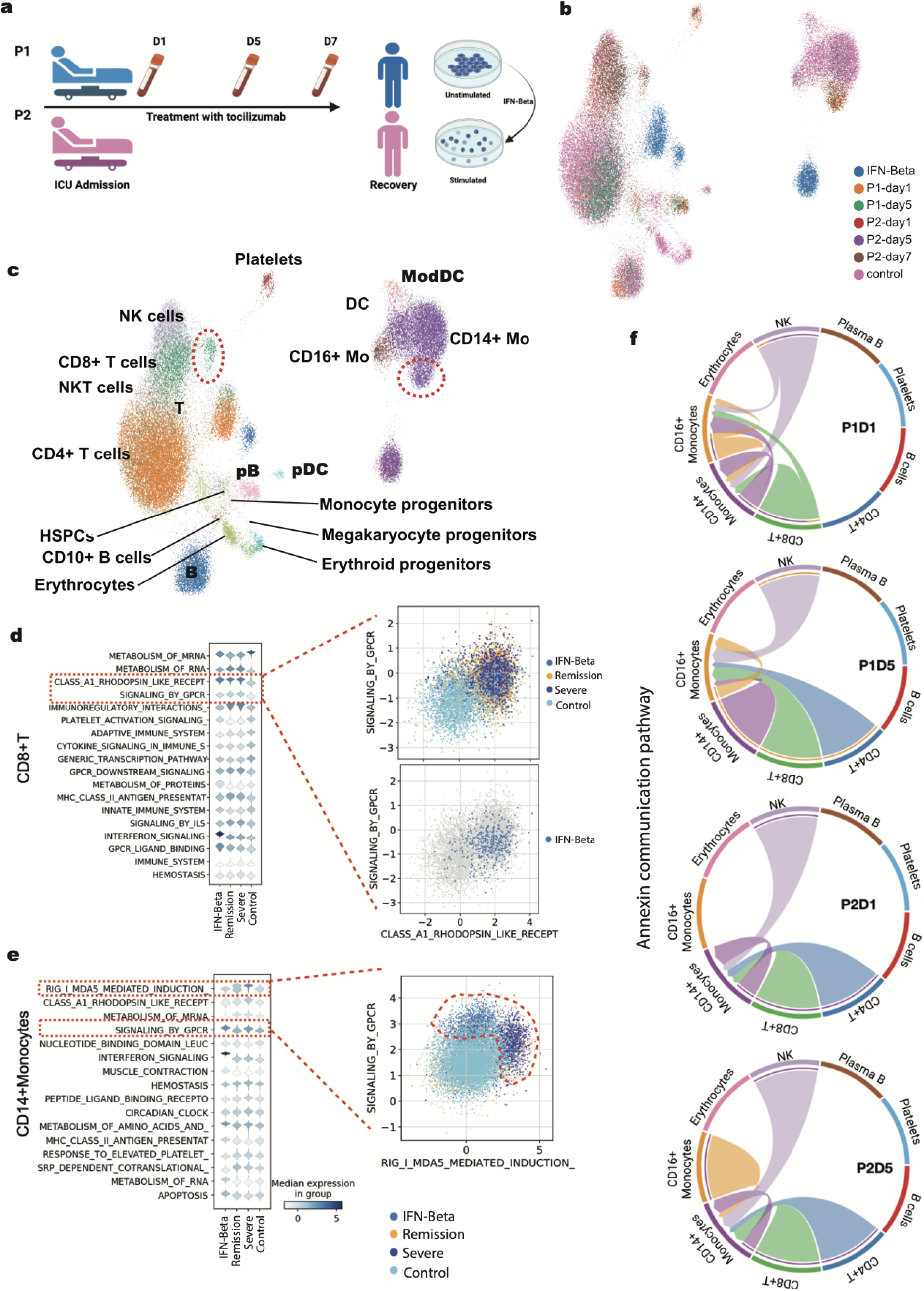
ExpiMap analysis highlights the importance of the annexin gene family communication pathway during moderate and severe COVID-19. **(a)**Illustration of the integrated datasets from PBMCs of healthy controls, patients with severe COVID treated with tocilizumab, and in the remission stages, and *in vitro* IFN-stimulated PBMCs. Figure made with Biorender. **(b)**Integrated manifold using expiMap showing combined healthy PBMCs (n = 32,484), two query datasets including two patients with COVID-19 (n = 18,752) and the IFN-β dataset (n = 13,576) from Kang *et al.* **(c)**Detailed cell type annotation of the integrated PBMC datasets. Red circles highlight cells that were not merged with the healthy PBMC cell atlas. DC = Dendritic cells, ModDC = Monocyte-derived dendritic cells, CD14^+^ Mo = CD14^+^ monocytes, CD16^+^ Mo = CD16^+^ monocytes, pDC = Plasmacytoid dendritic cells, pB = Plasma B cells. **(d,e)**Distribution plots for differential GP activities obtained using expiMap for CD8^+^ T cells and CD14^+^ monocytes, highlighting the antiviral transcriptional programs for *RIG-I/MDA5* and *GPCRs* in each population. Scatter Plot are latent GPs representation of highlighted GPs for each cell type. **(f)**Annexin communication pathways in different stages of COVID. In the severe stage (P1D1), both CD14^+^ and CD16^+^ monocytes participate in a dynamic communication circuit via annexins with NK and CD8^+^ T cells. This circuit converges to focused signaling to CD16^+^ monocytes during COVID remission (P1D5). In P2, CD14^+^ monocytes receive focused annexin signaling from NK, CD8^+^, and CD4^+^ T cells in the severe stage (P2D1), and later converges to signaling to CD14^+^ monocytes from the same lymphoid effectors during remission (P2D5).

Visual inspection of the joint representation of the query and reference pointed us toward the population of CD8^+^ T cells and monocytes (**Fig. 5c**) in both severe and remission stages of both patients that did not integrate into the healthy reference, unlike other populations from the same patients. We investigated this by performing a differential GP test between severe query cells and control cells in the reference for these cell types, with the aim of identifying GPs that could explain this separation. We identified transcriptional programs of antiviral response at different clinical stages of COVID-19 and in specific PBMC cell types. Pathogen recognition receptor (PRR) *RIG-I/MDA5* and GPCR pathways were shown to display differential behavior in CD8^+^ T cells (**Fig. 5d**) and CD14^+^ monocytes (**Fig. 5e**) in severe COVID-19 (D1) and during remission (D5, D7). The *RIG-I/MDA5* and GPCR pathways are known to initiate the innate immune response and modulate the adaptive immune responses during viral infections^57^; specifically, they are reported to coordinate the inflammatory response and its progression during COVID-19^58,59^. These findings suggest that a complex circuit of cellular communication may be differentially activated in both patients and may be related to the differences in treatment response at the cellular level.

To investigate this effect further, we estimated underlying cellular communication circuits using CellChat^60^, and compared them at different clinical stages in our integrated immune atlas. This analysis revealed that the annexin pathway displayed differential behavior in the severe and remission stages of P1 and P2, involving CD14^+^ and CD16^+^ monocytes, NK cells, and CD8^+^ T cells (**Fig. 5f** and **Supplementary Fig. 11**).Annexins are structural proteins that participate in multiple processes, including the regulation of inflammatory responses and homeostasis^61^; recently, they have been associated with the disease severity of COVID-19 in patients requiring hospitalization^62,63^. In this circuit, CD14^+^ and CD16^+^ monocytes receive signals from NK cells and CD8^+^ T cells for P1D1, whereas for P1D5, the annexin circuit switches completely to signaling between CD16^+^ monocytes and CD4^+^ T cells. In stark contrast, P2D1 is characterized by the annexin circuit focusing on CD14^+^ monocytes, which continues throughout the remission stage (P2D5), with the addition of CD16^+^ monocytes persisting toward D7 of remission (P2D7) (**Supplementary Fig. 11**).

Although expression levels of annexins have been described as biomarkers for the prediction of disease severity^63^, our analysis using expiMap is the first to describe the annexin circuit at the cellular level between monocytes (CD14^+^ and CD16^+^), NK cells, and T cells in COVID. The differences observed between patients in the expression of annexin-related interaction circuits may be related to the capability of viral clearance in each patient^56^ and the early expression of *FPR1* by CD16^+^ monocytes, which is associated with the early detection of pathogenic molecules and tissue damage^64^. Interestingly, our analysis shows the expression of *IFNG* for P2D7 by NK and CD8^+^ T cells (**Supplementary Fig. 12**), which may indicate a more complex antiviral response than in P1 that is independent of the symptomatic resolution attained by tocilizumab. Moreover, when contrasting our results with the annexin circuit in the data from IFN-stimulated cells, we observed that the cell–cell interactions using the annexin pathway were dominated by the expression of *ANXA1* in dendritic cells (DC) rather than *FPR1* in CD14^+^ monocytes (**Supplementary Fig. 12**).Our results do not illustrate the same circuit; however, this may indicate a lung-specific interaction operating in the lungs after the monocytes migrate to the lung tissue.

Although both these patients recovered after treatment with tocilizumab, clinical studies clearly demonstrate that this behavior is not consistent, and other factors, such as tocilizumab posology, may affect the clinical outcome^65^. At the cellular level, expiMap identifies transcriptional and cell–cell interaction circuits with the potential to be druggable, such as RIG-I/MDA5 and annexins, to help suppress cytokine storm syndrome in patients with COVID-19, which results in hospitalization.

### expiMap delineates pancreatic cell and subtypes after disease perturbation

As a final use case we asked if expiMap could assist with interpretable cell type annotation and the analysis of cell state heterogeneity. As example we used multiple scRNA-seq data sets from mouse pancreatic islets to capture beta cell heterogeneity, including data from young healthy mice on postnatal day 16 (dataset name: Fltp_P16)^66^, healthy young mice from a non-obese diabetic model before the onset of type 1 diabetes (T1D) at 5 weeks of age (dataset name: NOD)^67^, healthy adult mice and those exposed to chemically-induced stress (dataset name: spikein_drug)^68^, and healthy adult mice and those exposed to chemically-induced (streptozotocin-induced) type 2 diabetes (T2D) with and without therapy with different combinations of insulin, GLP1, and estrogen (dataset name: STZ)^69^.

We used expiMap to integrate three non-T2D pancreatic datasets differing in multiple biological factors, including sex, age, and stress status (Fltp_P16, NOD, and spikein_drug, from here on named as the reference datasets) using PanglaoDB marker sets and Reactome pathways. PanglaoDB marker sets were used to enable cell type identification, as previously proposed for scRNA-seq annotation^70,71^. As an alternative, we used Reactome pathways to enable the identification of molecular processes^72^ differentially active across biological conditions. We projected one dataset (STZ, from here on named as the query dataset) that included healthy and T2D cells onto this reference using the architecture surgery approach (**Fig. 6a,b**).On the integrated embedding (**Fig. 6b**), a residual separation between studies is observed, as is expected owing to biological differences between the integrated mice models, such as disease state and age. To evaluate expiMap integration performance, we also performed integrations with scArches +scVI, Seurat V4, and Symphony (**Supplementary Fig. 13a**) and assessed the integration quality using scIB metrics (**Supplementary Fig. 13b**);our findings showed that, for all metrics, expiMap performs similarly to existing methods or outperforms them.

**Fig. 6.**
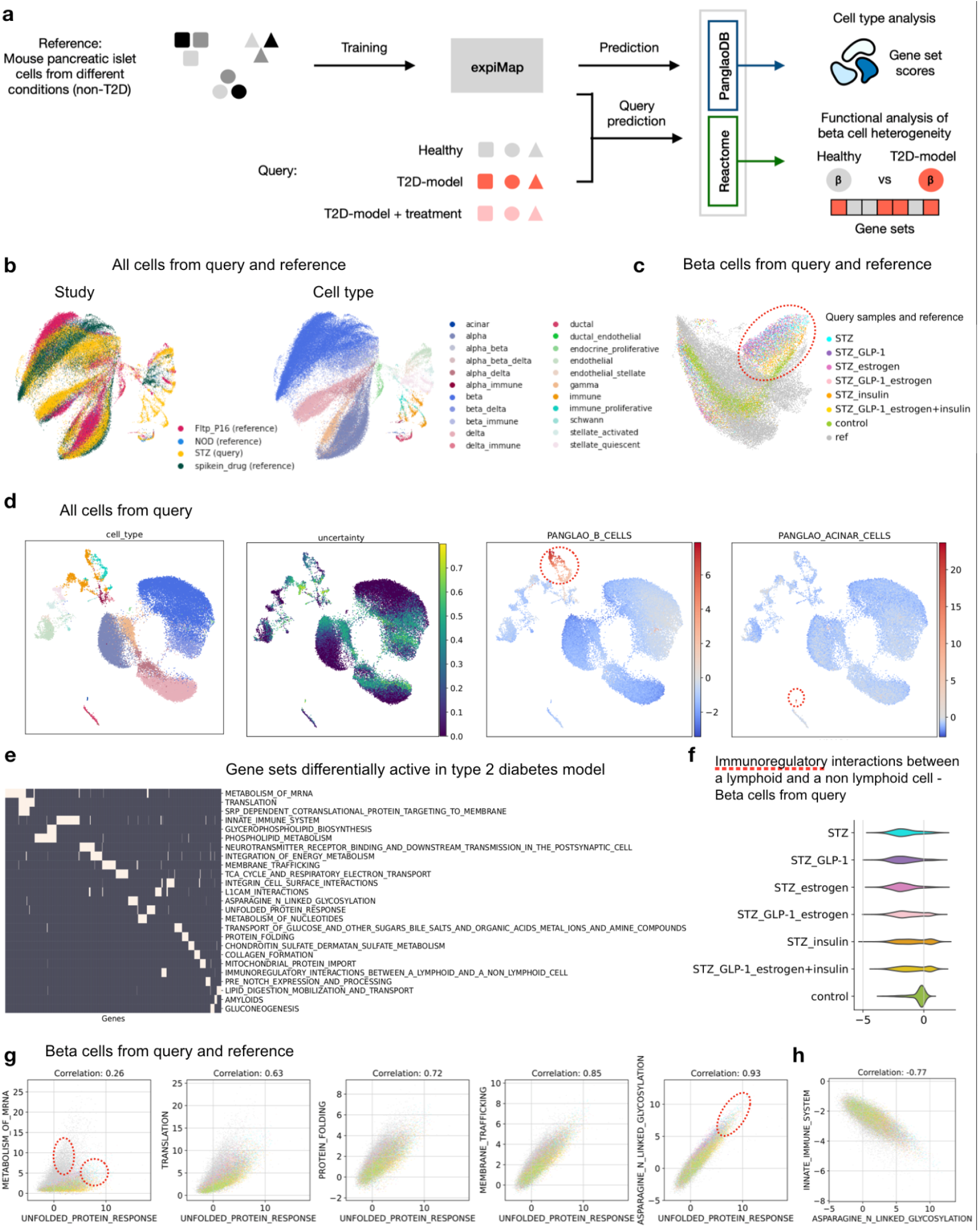
Reference mapping of pancreatic islet cells using expiMap. **(a)**Process of the pancreatic islet cell analysis. We trained the expiMap model on a heterogeneous set of non-T2D mouse pancreatic islet cells from different studies with GPs from PanglaoDB and Reactome. A dataset containing healthy, T2D-model, and treated T2D-model cells was then mapped to this reference. expiMap scores from PanglaoDB helped us to evaluate cell type annotation and scores from Reactome to determine metabolic differences between healthy beta cells and T2D-model beta cells. **(b)**UMAP colored by dataset shows three reference pancreas datasets (45,178 cells) and one query dataset (36,899 cells) integrated with expiMap. **(c)**UMAP of beta cells separates healthy beta cells and T2D-model beta cells from the reference and query datasets. **(d)**expiMap scores help with manual cell type annotation. The score for immune B cells highlights a subpopulation of cells that was previously annotated under the umbrella term of immune cells. The score for acinar cells helps in the annotation of the small acinar cell type cluster, which was not annotated in automatic cell type transfer owing to low classifier certainty (rightmost plot). **(e)**Low redundancy of the top differential Reactome pathways between healthy and T2D-model query beta cells, shown by the low overlap between pathways. Genes (columns) associated with each GP (rows) are marked in white; the absence of a gene in a GP is indicated by dark gray color. Genes and GPs were clustered based on the displayed gene–GP association matrix. **(f)**The immune interaction GP scores in insulin-treated T2D-model beta cells from the query exhibit bimodal distribution. **(g,h)**Beta cell scores of selected GPs differentially active in T2D-model beta cells enable the comparison between per-sample distributions and the identification of GPs that are highly correlated across all datasets. Legend is shown in panel **c.**On the first subplot, circles mark the T2D-model population with relatively high scores in UPR and mRNA metabolism compared with the healthy control from the query, whereas other non-T2D cells from the reference show high mRNA metabolism without a high UPR score. On the final subplot, the circle indicates the T2D-model population with extreme UPR and asparagine *N*-linked glycosylation scores. ref: reference datasets, other samples are from the query dataset; STZ: streptozotocin T2D-model; STZ_GLP-1: streptozotocin T2D-model treated with GLP-1; STZ_estrogen: streptozotocin T2D-model treated with estrogen; STZ_GLP-1_estrogen: streptozotocin T2D-model treated with GLP-1-estrogen conjugate; STZ_insulin: streptozotocin T2D-model treated with insulin; STZ_GLP-1_estrogen+insulin: streptozotocin T2D-model treated with GLP-1-estrogen conjugate and insulin; control: healthy control.

Next, we automatically transferred cell type annotations from reference to query, (**Fig. 6d, Supplementary Fig. 14**).Analyzing expiMap-generated scores of pancreatic cell type-associated PanglaoDB GPs (**Supplementary Fig. 14c**) helped with the annotation of ambiguous cell clusters (**Supplementary Fig. 14b**): For example, expiMap scores helped to resolve potential doublets (e.g., immune–endocrine doublets) and small cell populations (e.g., acinar cells) that were marked as unknown or wrongly annotated (**Fig. 6d, Supplementary Fig. 14b,c**).This is relevant as automated cell type annotation methods often produce unreliable results in challenging cell populations, such as doublets, rare cell types, or transitional cell states, and require manual assessment. It is usually done by plotting marker expression, which requires corrected data for comparability^70^, while expiMap scores are directly comparable. Similarly, expiMap scores can resolve coarsely-annotated cell types; in our case the B cells that we annotated under the joint term of immune cells (**Fig. 6d**).Furthermore, expiMap integration preserves cell subtype variability. Namely, expiMap-based UMAP of beta cells shows gradual separation of diseased, differently treated, and healthy query cells (**Fig. 6c**) according to known treatment efficacy: GLP-1-estrogen+insulin was the most effective^69^. Healthy beta cells from the reference and query exhibit greater overlap. The same pattern is quantitatively supported by the PAGA graph, which indicated the strongest connection of reference cells with the query control, followed by insulin-treated T2D-model samples, and weaker connections with other T2D-model samples (**Supplementary Fig. 13c**).To evaluate if expiMap has captured the diabetes-associated beta cell loss of identity, dedifferentiation, and transdifferentiation to other endocrine cell types^69,73^, we investigated the relevant cell type GPs (**Supplementary Fig. 13d**).Indeed, in T2D-model beta cells, we observed lower scores for beta cell identity markers in conjunction with higher scores for markers of pancreatic progenitors and enteroendocrine cells (an umbrella term that also encompasses pancreatic endocrine cells). This shows that expiMap can capture heterogeneity within relatively homogenous populations, such as a particular cell type, and thus helps us to interpret why batches do not fully overlap within the embedding. Our analysis thus indicates that expiMap produces high-quality integrations, which can be directly interpreted at the molecular level.

To search for molecular changes between the healthy control and T2D-model beta cells from the STZ study, we used the expiMap enrichment function with the Reactome GP collection (**Supplementary Table 2**).As we had set strong regularization for Reactome GPs, we observed little overlap between the enriched GPs (**Fig. 6e**), simplifying the interpretation of enriched GPs. We observed differences in energy metabolism and protein synthesis, the unfolded protein response (UPR), cell–matrix interactions, and cell–cell interactions, including Notch signaling and immune communication. Genes that most strongly contributed to the activation of the enriched GP are reported in the **Supplementary Table 3**. Some of these differences were already reported in the original study, namely the diabetes-associated increase in oxidative phosphorylation, electron transport chain activity, and endoplasmic reticulum stress, which is related to the UPR^69^. The remaining identified differences were not highlighted in the original study, although other studies observed diabetes-related increase in insulin protein synthesis^74^, Notch signaling^75^, disrupted islet architecture^76^, and immune infiltration into islets^77^. Together, these results demonstrate that expiMap enrichment was able to identify diabetes-associated molecular changes in the T2D-model cells.

As expiMap computes GP scores for each cell, we performed a cross-sample comparison of the distributions of Reactome GP scores differentially active between the T2D-model and healthy beta cells. We find that interactions between lymphoid and non-immune cells (**Fig. 6f**) have a multimodal distribution within T2D-model beta cells treated with insulin, potentially indicating the presence of multiple cell states within individual samples. For scores from other enriched GPs, we did not observe such distinct multimodal patterns within individual samples. As expiMap score distributions capture multimodal patterns, there may be a need for the use of specialized differential testing methods that can account for multimodality^78,79^.

The UPR arises in T2D owing to the proinsulin synthesis rate, which exceeds the protein processing capacity of cells, leading to beta cell dysfunction and death^80,81^. Thus, we attempted further study of the enriched GP describing the UPR, and compared the distribution of UPR scores with the scores of other GPs associated with protein synthesis and processing (**Fig. 6g**).As expiMap produces batch-corrected GP scores across all cells, we also plotted the distribution of these scores in the reference, enabling a direct cross-study comparison. We observed a high correlation between the UPR and asparagine *N*-linked glycosylation GP scores (absolute correlation coefficient of 0.93) across all datasets with extreme scores in T2D-model cells (**Fig. 6g**).An increase in *N*-linked glycosylation has been previously implicated in diabetes, although the regulatory background is not clear^82,83^. The observed correlation between the UPR and *N*-linked glycosylation across a range of different beta cell states from different studies may implicate a shared regulatory mechanism between the UPR and *N*-linked glycosylation in beta cells. Indeed, it has been previously shown that the UPR regulator XBP1 affects *N*-glycan structures^84^; however, to the best of our knowledge, we are the first to report this correlation in pancreatic beta cells. It was suggested that these changes in *N*-glycans on the cell surface may be also involved in cellular signaling and the immune response^84^. Thus, we checked if immune-related GPs differentially active in T2D-model beta cells correlate with *N*-linked glycosylation, as it is known that immune infiltration plays a role in T2D and may be caused by metabolic stress and changed epitopes on beta cells^77,85^. Again, we observed a strong correlation between *N*-linked glycosylation and the innate immune system GP scores (absolute correlation coefficient of 0.77) across datasets (**Fig. 6h**), but the correlation with other GPs was lower (**Supplementary Fig. 15**).We believe that further research into connections between these GPs may reveal new knowledge regarding the metabolic stress response in beta cells. Collectively, these results demonstrate that the analysis of expiMap GP activity distributions provides insights beyond simple differential activity analysis.

## Discussion

We introduced expiMap for interpretable single-cell reference mapping. Our model embeds domain knowledge in the form of gene programs (GPs) into the deep learning architectures used for reference mapping, and is able to complement these GPs further with newly discovered unconstrained GPs for query data sets. The interpretability of the model allows the users to generate immediate inferences about the query once mapped to a reference within the context of GPs. This is in contrast to the existing analysis pipelines, which involve multiple steps and without end-to-end learning necessarily aggregate processing errors from previous steps. Interestingly, in a comparison across five different organ atlases, we found that the constrained expiMap model did not lose expressiveness versus an unconstrained CVAE model; indeed, prior constraints appeared to improve reference mapping and *de novo* data integration performance, confirming the earlier concepts of adding “differentiable programs”^19^. Through various applications, we demonstrated the interpretability of the model. Our analysis of interferon response at the single-cell level revealed the GPs involved in cell type-specific responses to IFN perturbation. The explainability of the model also allowed us to determine why specific cell types from patients with COVID-19 did not respond to an immunosuppressive drug prescribed to prevent potential cytokine storm. Finally, we leveraged expiMap to achieve accurate annotation of new query pancreatic cells and, further, analyze the heterogeneity in pancreatic beta cell states.

Reference mapping with expiMap provides a new perspective to data integration and reference mapping. In scenarios with large differences in the datasets, such as cross-species mapping, the query data might not be fully aligned in the reference owing to the substantial biological and technical differences dominating the overall representation obtained by existing methods. This phenomenon makes it challenging to distinguish shared and unique signals between datasets. expiMap enables the integration of datasets along the axes of variations explained by a single or multiple GPs, where the datasets share variations and are mixed. This mixing stems from the commonality of the datasets in those programs. Such insights could not be obtained by, for example, looking at the overall UMAP, which would be influenced by all genes, might be misleading and could obscure such commonalities. As we demonstrated when mapping COVID-19 patient data, CD8^+^ T cells from patients with COVID-19 were separate from IFN-β-treated CD8^+^ T cells in the global representation obtained from all GPs in UMAP (see **Fig. 5b, c**). At the same time, they are integrated within specific GPs, capturing shared signals in two different cell states (see **Fig. 5d**). Overall, expiMap can provide more insights into data integration by contextualizing it within GPs. We also performed an evaluation of integration performance.

The idea of constraining VAE decoding is not new. We first proposed a regularized linear decoder to include domain knowledge into autoencoders for single-cell data at a conference^86^, with scalable and expressive embeddings when compared to existing factor models, such as f-scLVM^87^. Recent approaches such as VEGA^88^, scTEM^89^ and pmVAE^90^ also feature VAE-based architectures with linear decoders or training separate VAEs for each GP yet connected via a global loss in the case of pmVAE. In contrast, expiMap aims toward interpretable reference mapping allowing to fuse reference atlases with GPs and enabling the query of genes or GPs. Overall, we see the novelty of expiMap in its ability to concurrently encode domain knowledge to enrich the existing GPs with new information while refining existing knowledge and learning new knowledge *de novo* from the data for contextualized integration of query datasets into references. Finally, the usage of proximal optimization and operators (see **methods**) in expiMap is theoretically different from previous methods, which is crucial for the joint soft membership of genes and GP selection using the group lasso feature.

Our model leverages domain knowledge to improve the interpretability of deep learning models useful for single-cell genomics. With increasing availability ^91,92^ of curated domain knowledge, expiMap can be trained on multiple databases while pruning irrelevant information. However, selecting the relevant knowledge to include in the model can affect the model’s performance. As we demonstrated, including interferon-related knowledge can improve the performance in reference mapping (see **Fig. 2**), while excluding it can lead to poor mapping of the query (see **Supplementary Fig. 6**). Another limitation concerns the interpretation and validation of newly learned GPs that capture new variations in the query data. As we demonstrated, looking at distribution plots and visualizing the embedding can decipher the variations. However, the validation at the gene level requires further expert knowledge for each biological system. Another limitation is the modeling hierarchies in unsupervised settings, starting from single genes to GPs and to higher-level biological processes. Previous work, such as knowledge-primed neural networks (KPNNs)^93^, P-net^25^ and visible neural networks (VNNs)^94^ employed hierarchical modeling, but in supervised settings, to predict tumor type or cell states. Using a similar strategy in an unsupervised model would add another layer of analysis to mapping data, not only to GPs but also to biological processes, and potentially improve the performance. A final limitation of general DL models may be data hunger. To determine the sensitivity of our model to dataset size, we trained models of increasing quality by incrementally including more training samples in the reference building task (see **Supplementary Fig. 16**). We observed that expiMap outperformed the linear baseline of a non-biologically informed linear decoder model (LDVAE) in a low-data regime. The more complex non-amortized scVI achieved the best results with increased number of training samples, while expiMap outperformed scVI and LDVAE. Overall, these results suggest that incorporating prior knowledge leads to more sample-efficient learning in the presence of fewer samples than non-biologically informed models with similar complexity (e.g., LDVAE). Further, when more training samples are available to learn GP activities efficiently, expiMap performs with complex non-linear models.

Although we demonstrated expiMap by using scRNA-seq data, the model is naturally extendable to multimodal^95,96,97^ datasets. Recent technological advances in single-cell biology allow the simultaneous capture of chromatin accessibility, gene expression, and protein levels in single cells^4^. This makes it possible to learn the hierarchy of connected representations by distilling domain knowledge about regulatory elements, transcription, and translation, covering multiple cellular processes into the representation learning methods. Another potentially exciting direction is the combination of the expiMap architecture with *in vitro* perturbation modeling approaches^5,6,27^ to model *in vitro* perturbations of GPs. Finally, given the availability of spatial transcriptomics data^98^, it is possible to adapt expiMap to include information about cell-to-cell communication^99^ in the learned representations to gain further insights into cellular communications and signaling.

Researchers in the field of single-cell genomics are moving toward using reference mapping to analyze new query datasets. One example is Azimuth^8^, a web tool that enables the users to map their data into a reference for visualization and annotation. We envision expiMap will further advance the applicability of reference mapping methods by bringing a new layer of interpretability and mechanistic understanding to integrative single-cell data analysis facilitating biological hypothesis generation and discovery.

## Data availability

All datasets used in the paper are public, referenced and downloadable at https://github.com/theislab/expiMap_reproducibility.

## Code availability

The software is available as part of https://scarches.readthedocs.io/en/latest/.

The code to reproduce the results is available at https://github.com/theislab/expiMap_reproducibility.

## Acknowledgments

We are grateful to all members of the Theis laboratory. M.L. is grateful for valuable feedback on the text from Fabiola Curion and Luke Zapia. M.L. is thankful for feedback from Adam Gayoso on amortized inference. M.L and K.H. acknowledge financial support from the Joachim Herz Stiftung via Add-on Fellowships for Interdisciplinary Life Science. K.H. acknowledges support from Helmholtz Association under the joint research school “Munich School for Data Science.” This work was supported by the BMBF (01IS18036A and 01IS18036B), by the European Union’s Horizon 2020 research and innovation program (grant 874656), by Helmholtz Association’s Initiative and Networking Fund through Helmholtz AI (ZT-I-PF-5-01), and sparse2big (ZT-I-0007), all to F.J.T.

## Author contributions

M.L. and S. R. conceived the project with contributions from F.J.T. and S.R. S.R and M.L. implemented and trained the models. M.L designed the experiments. M.L, C.T.L, K.H, and S.R performed the analysis. A.M helped to design the experiment related to COVID-19. S.H.Z performed the limma-fry comparison. F.J.T supervised the research. All authors wrote the manuscript.

## Competing interests

F.J.T. consults for Immunai Inc., Singularity Bio B.V., CytoReason Ltd, and Omniscope Ltd, and has ownership interest in Dermagnostix GmbH and Cellarity.

## Supplementary materials

**Supplementary Table 1:** List of genes learned by B cell node + DEG obtained from differential testing in Scanpy

**Supplementary Table 2:** Differential Reactome terms analysis between healthy control and T2D-model samples from STZ study. Bf: Bayes factor.

**Supplementary Table 3:** Gene weights for activation of terms reported in Supplementary Table 2. Genes are ordered from strongest to weakest term activation based on absolute weight. Genes with zero-weight in each GP are not reported. EID - Ensembl ID.

**Supplementary Table 4:** Information on pancreatic datasets used in this study.

**Supplementary Table 5–16**: Information about training and model hyperparatmers for all experiments reported in the paper.

## Supplementary figures

**Supplementary Figure 1.**
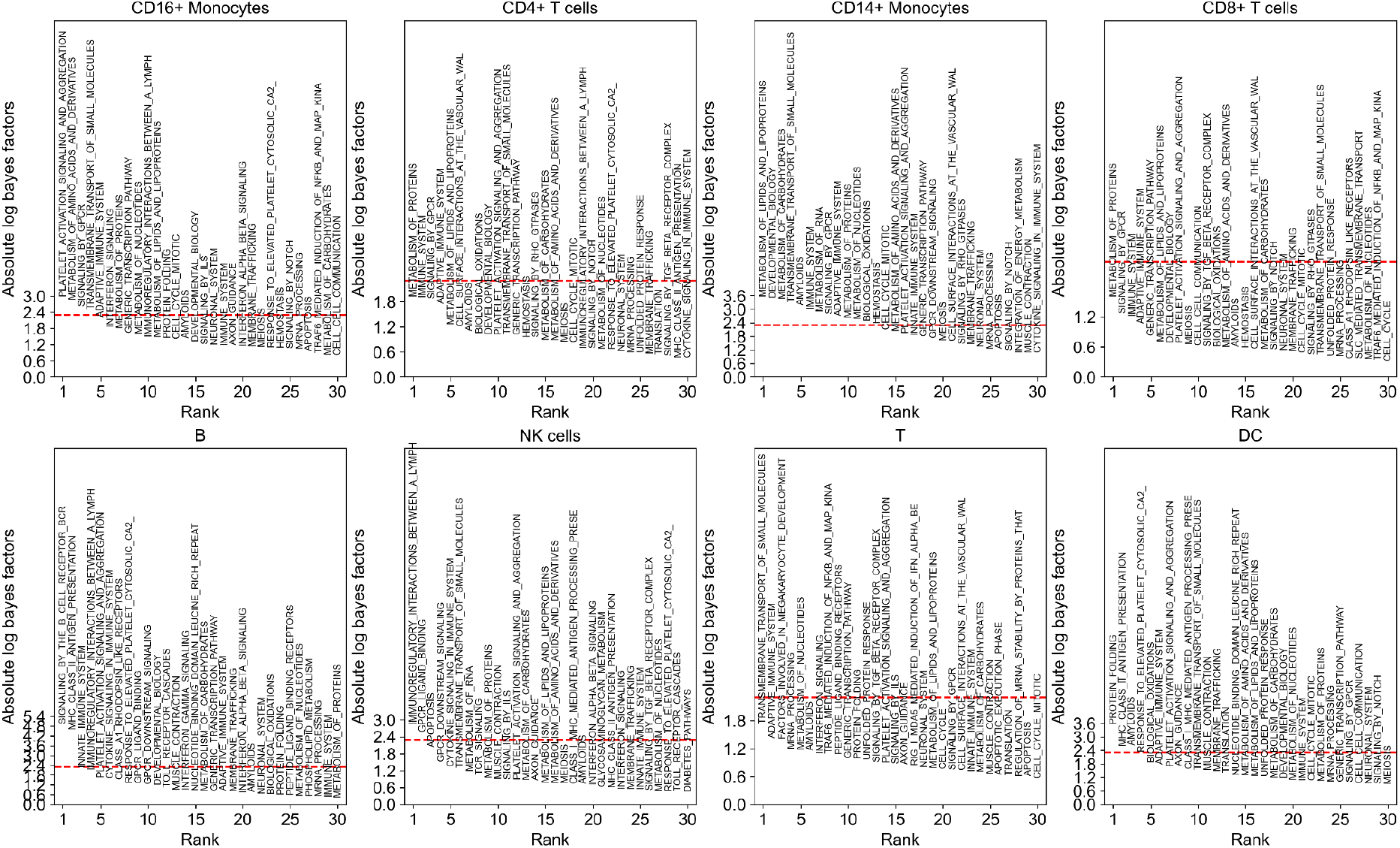
Differential GP analysis results for cell types from the integrated query and reference PBMCs with five datasets. Differential GP analysis results between cell types (one vs all test) for cell types in the query data. The x-axis is the ranking of GPs; the y-axis denotes the significance (absolute log-Bayes factor) of each GP.

**Supplementary Figure 2.**
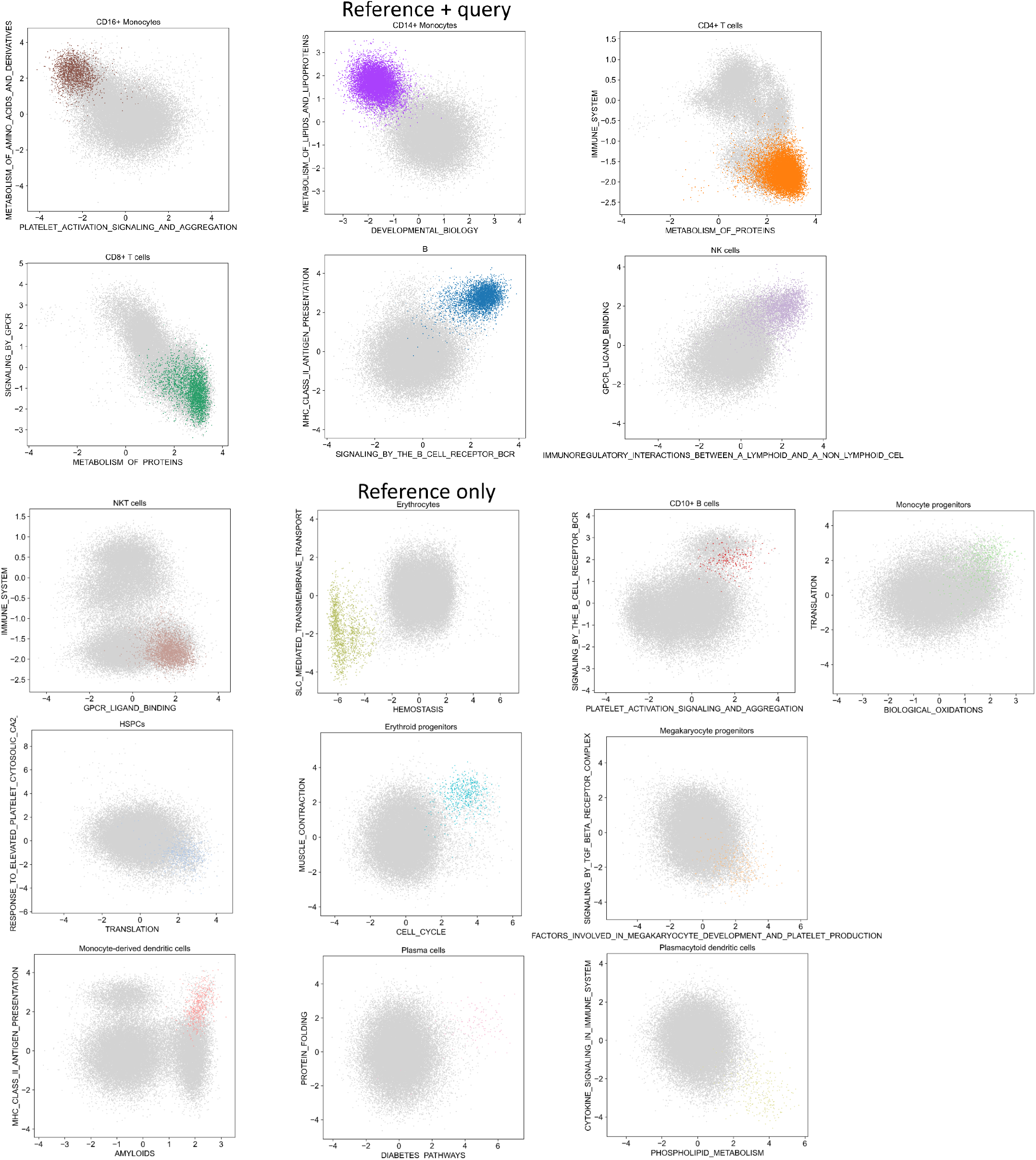
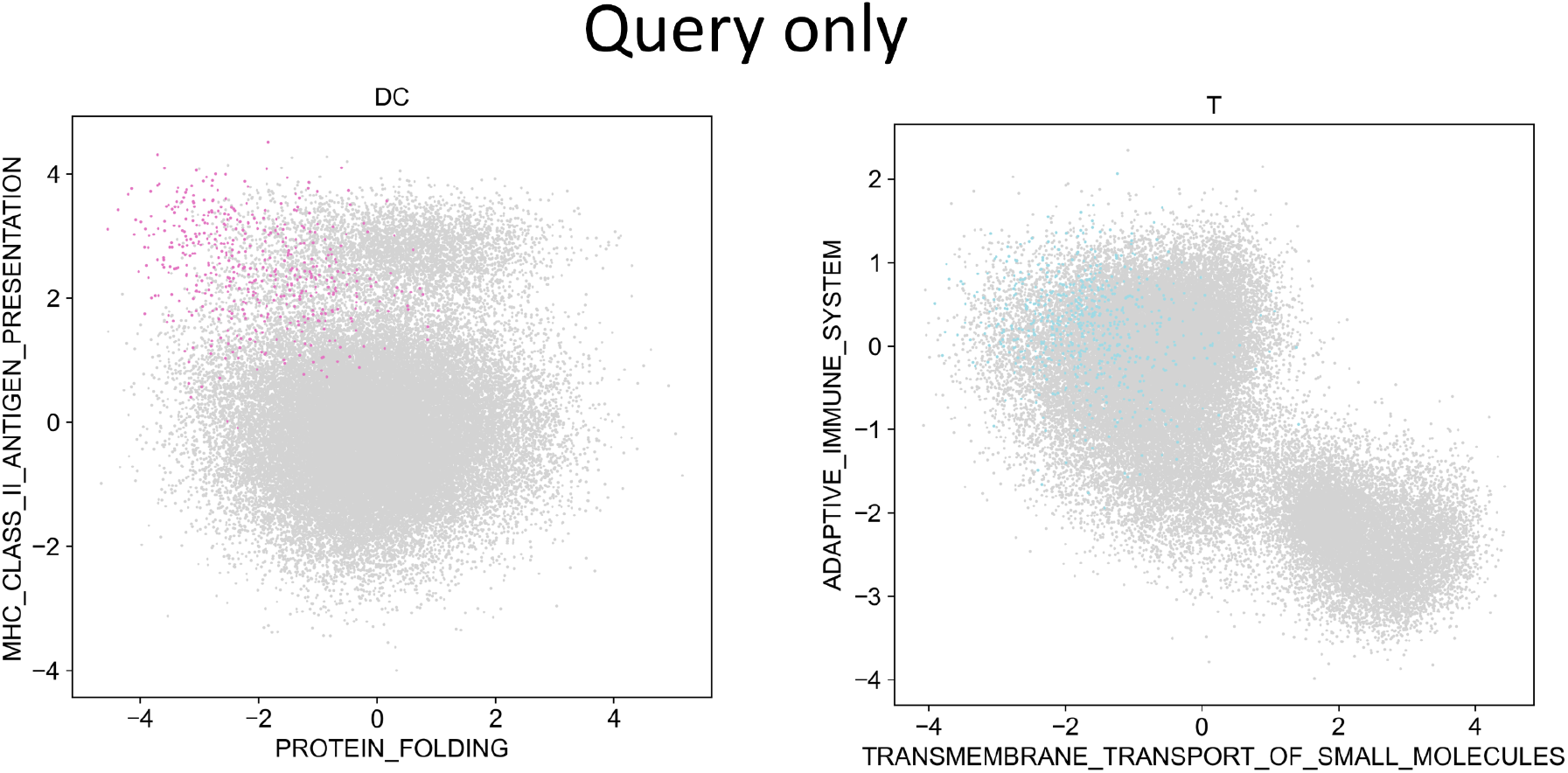
Differential GP analysis results for cell states. Two-dimensional visualization of the top two GPs resulting from the cell type differential analysis results for cell types in the query, the reference, and shared among both. The colors highlight the cell types in the title of each panel while other cells are colored gray.

**Supplementary Figure 3.**
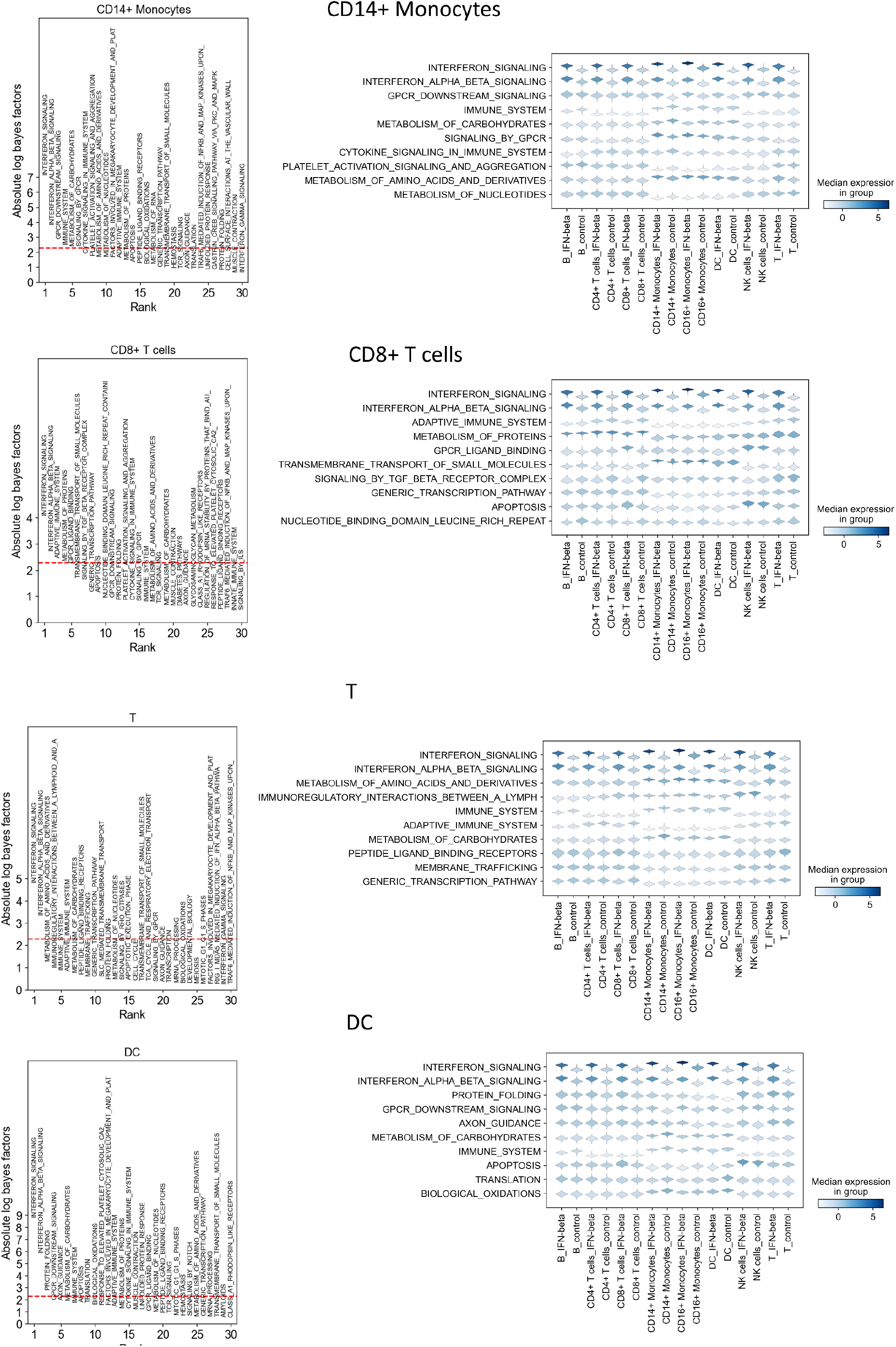
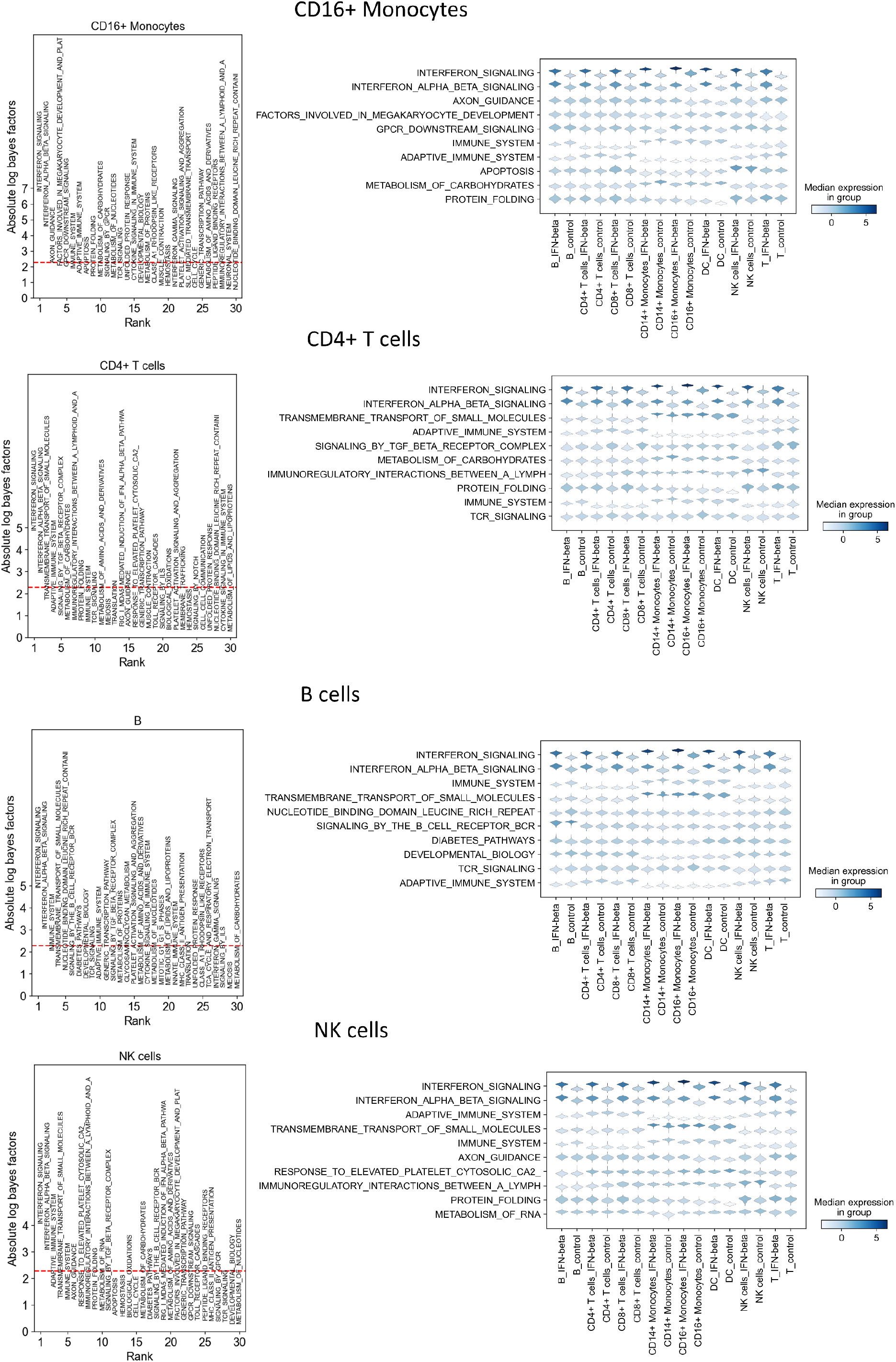
Cell type-specific differential GP analysis for cell states. Differential GP analysis results between IFN-β and control for cell types in the query data. The x-axis shows the ranking of GPs; the y-axis denotes the significance (absolute log-Bayes factor) of each GP. The violin plots demonstrate the activity of those GPs across other cell types.

**Supplementary Figure 4.**
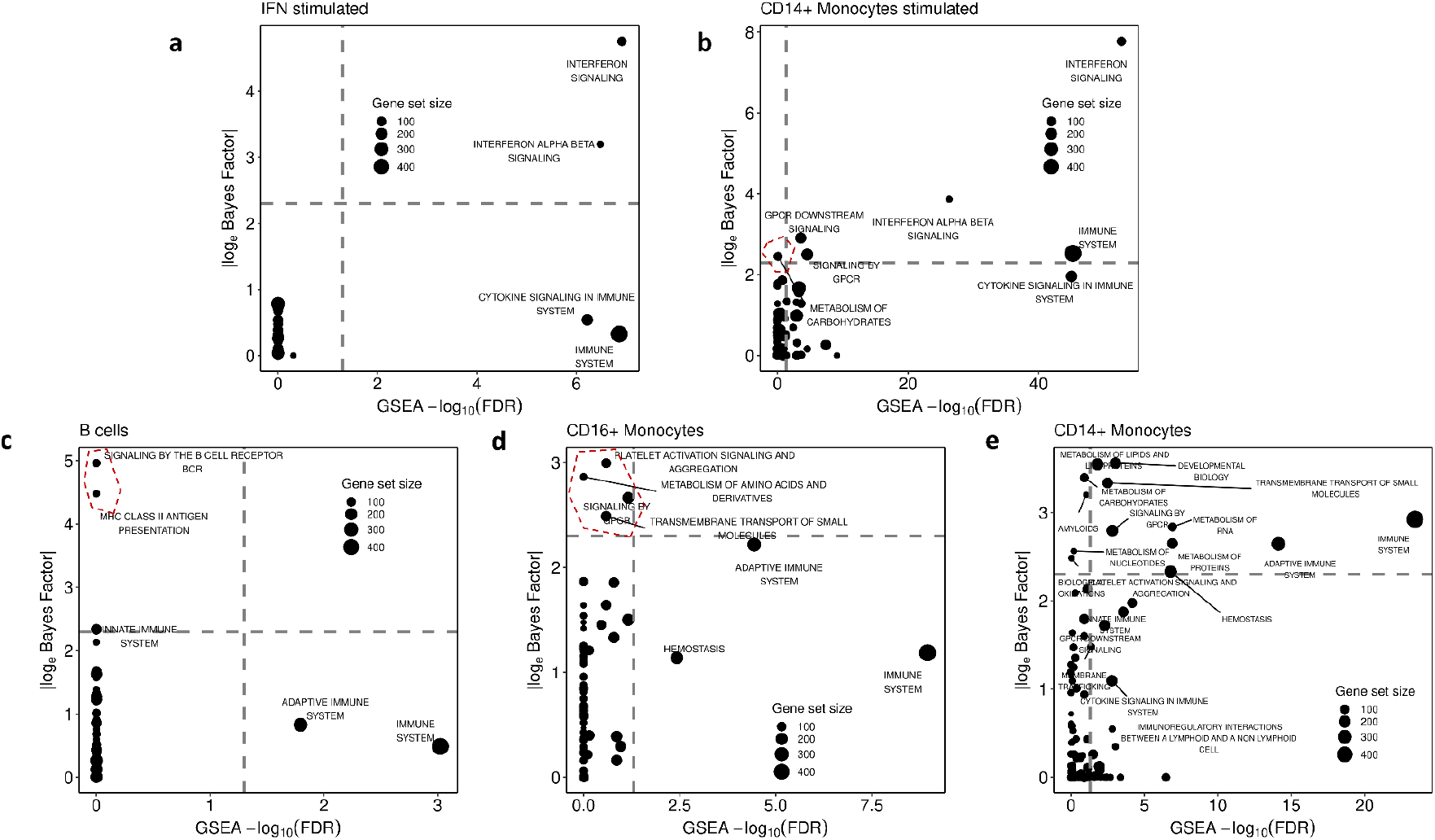
Comparison of expiMap Bayes Factor with GSEA-log10(FDR). For comparison, we show Bayes factors from expiMap and FDR from fry. **(a)**Overall stimulated vs control tests (all cell types are pooled together). **(b)**Results for CD14^+^ monocytes stimulated vs control tests. **(c)**B cells vs the rest of the cell types. **(d)**CD16^+^ monocytes vs the rest of the cell types, **(e)**CD14^+^ monocytes vs the rest of the cell types. The x-axis shows the negative logarithm of the false discovery rate (− *log*_10_ *FDR*) from fry; the y-axis shows the absolute value of the logarithm of the Bayes Factor (|*log*_e_(*Bayes Factor*)|) from the expiMap test. The size of the circles is proportional to the size of the gene set. We observed an overall agreement between expiMap and conventional GSEA results; however, expiMap detected more specific gene programs in some comparisons, while being computationally efficient with regard to runtime as the differential gene expression and enrichment testing steps no longer have to be repeated for every individual comparison.

**Supplementary Figure 5.**
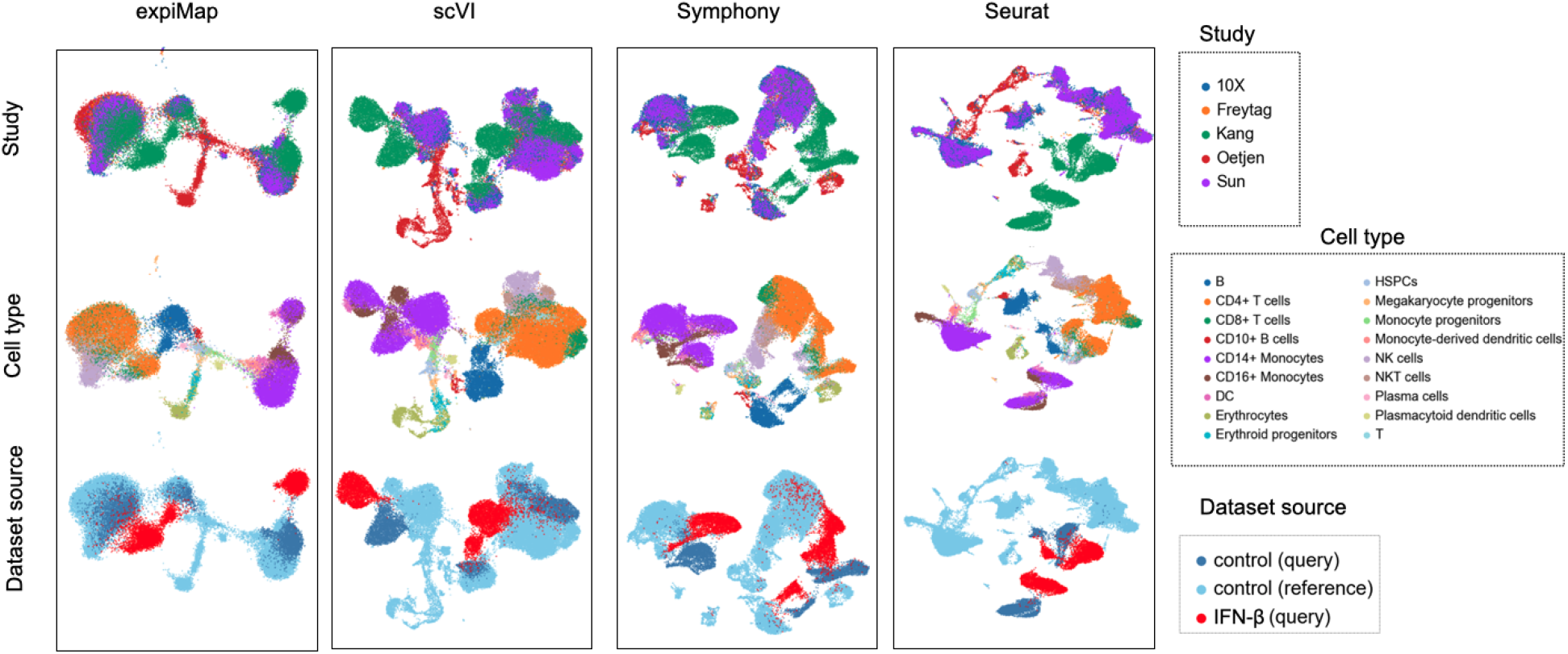
Benchmarking reference mapping methods. UMAP representation of integration accuracy of mapping IFN-β data onto the healthy atlas across studies, cell types, and dataset source for different models.

**Supplementary Figure 6.**
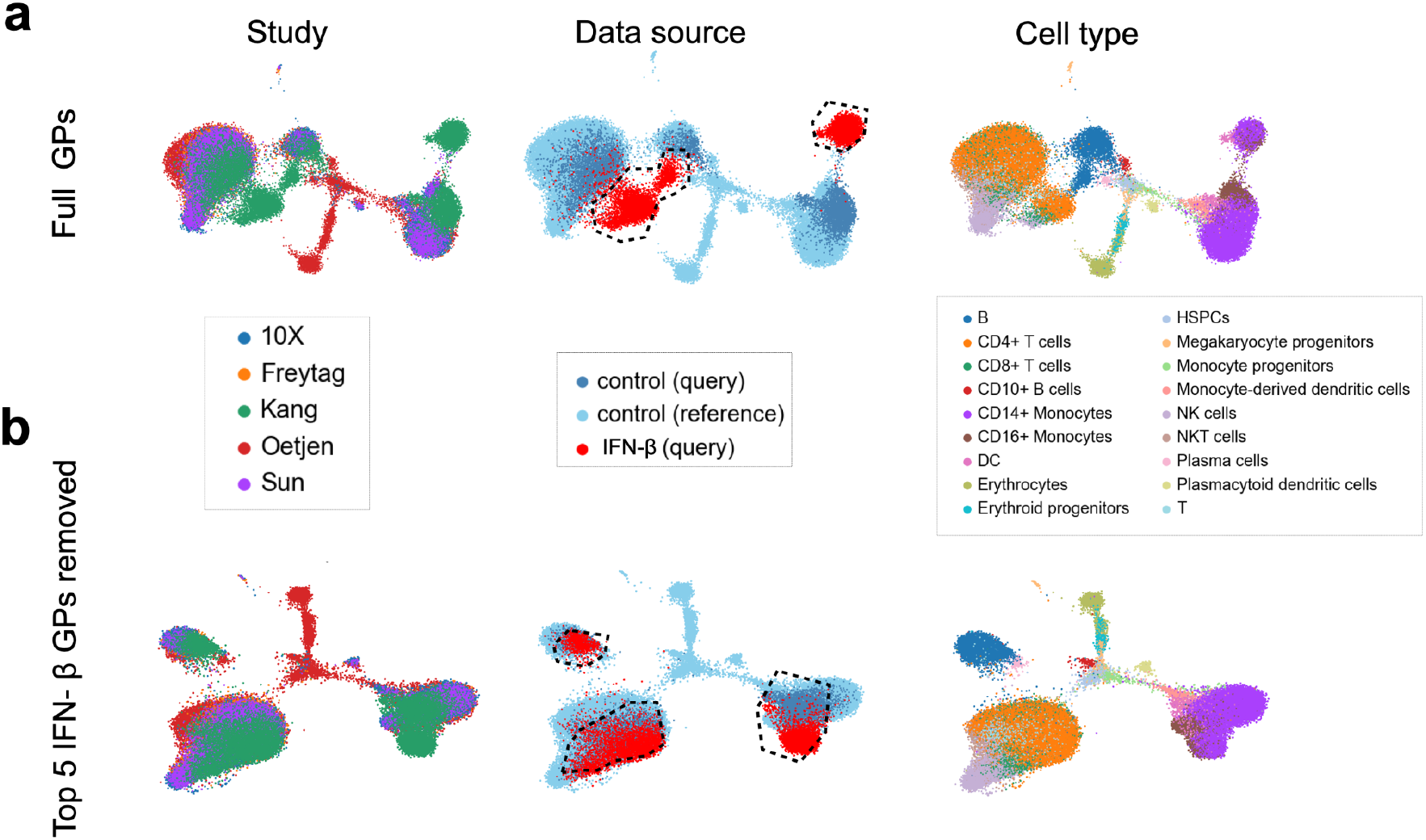
Assessing the quality of reference mapping by removing GPs. **(a)** Integration reference and query representation across studies, data source, and cell types when expiMap is trained with all available GPs. The highlighted populations are IFN-β-stimulated cells separated from the control samples. **(b)** Same model as **(a)** but the top five IFN-β stimulation-related pathways were removed from training and the model is unaware of them. The highlighted populations are IFN-β-stimulated cells that the model incorrectly merged into control cells, removing perturbation heterogeneity in the query data.

**Supplementary Figure 7.**
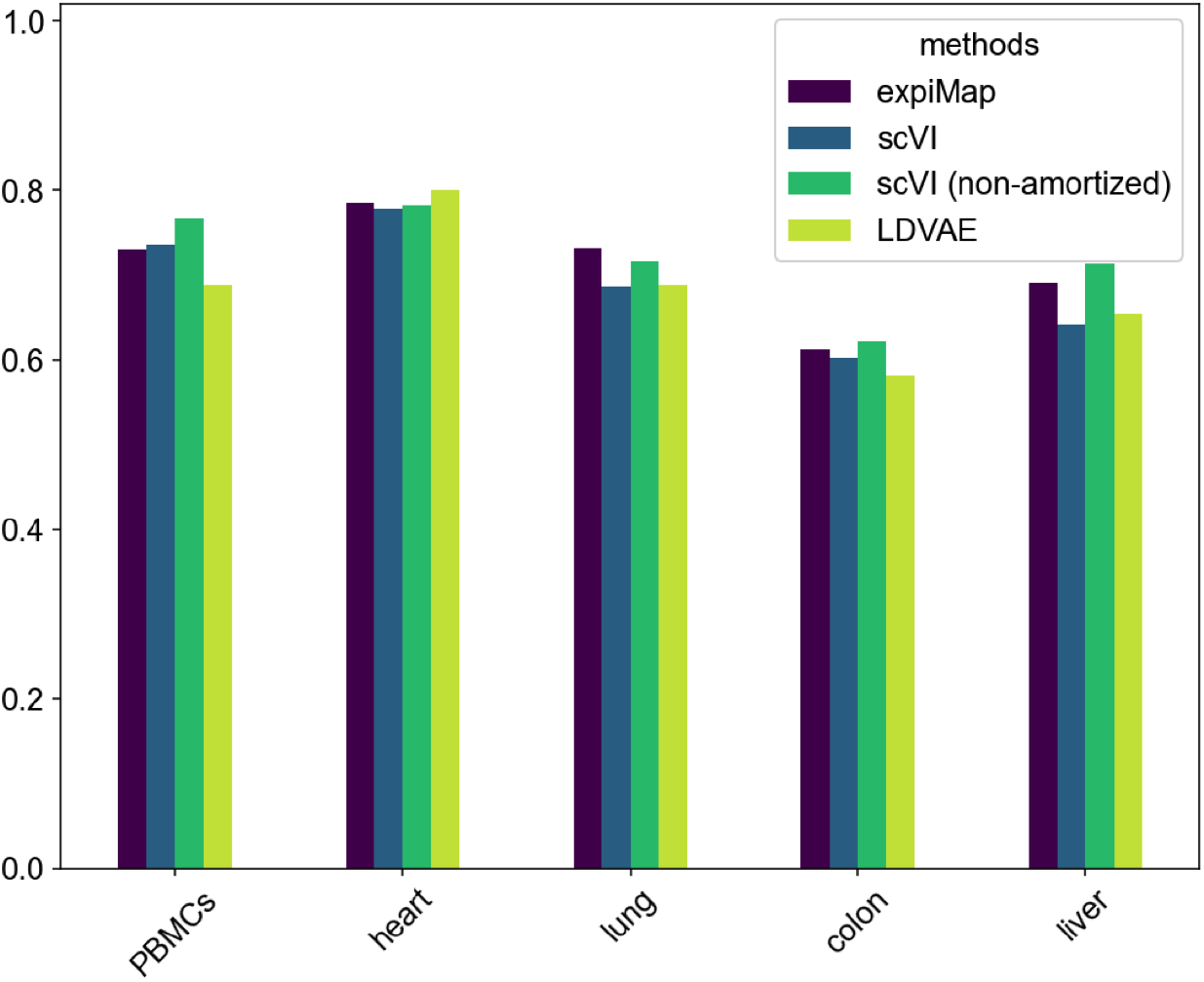
Benchmarking the reference building ability of expiMap. Comparison of the reference building performance by benchmarking across five different tissues, PBMCs (n = 161,764), heart (n = 18,641), lung (n = 65,662), colon (n = 34,772), and liver (n = 113,063), and four different methods.

**Supplementary Figure 8.**
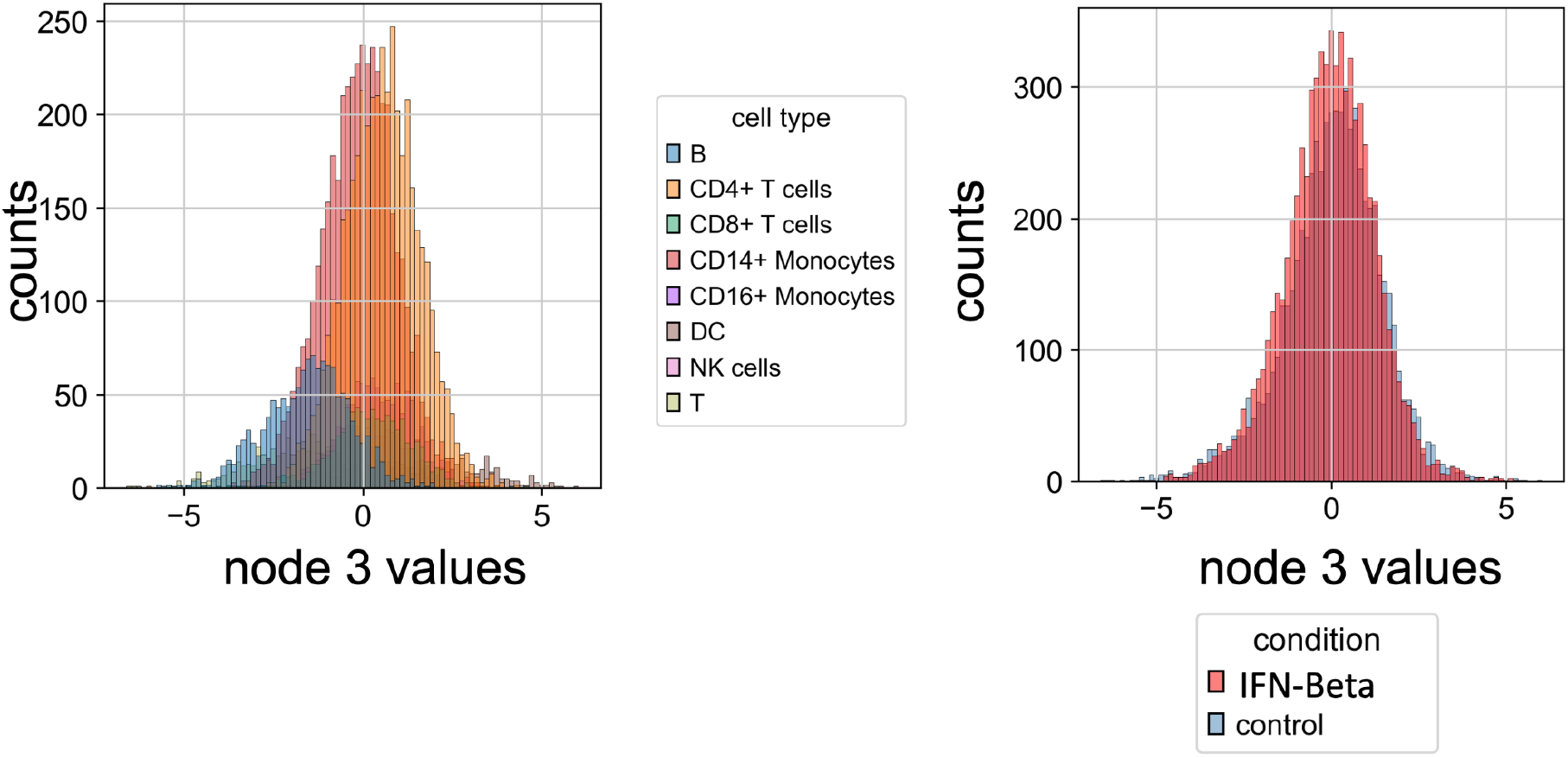
Activity distribution for an inactive program. Histogram of activity values colored by cell type (first column) and condition (second column) for a new GP learned during query training, demonstrating the lack of discriminative information for any specific cell type or condition. The x-axis represents the activity values for each single cell.

**Supplementary Figure 9.**
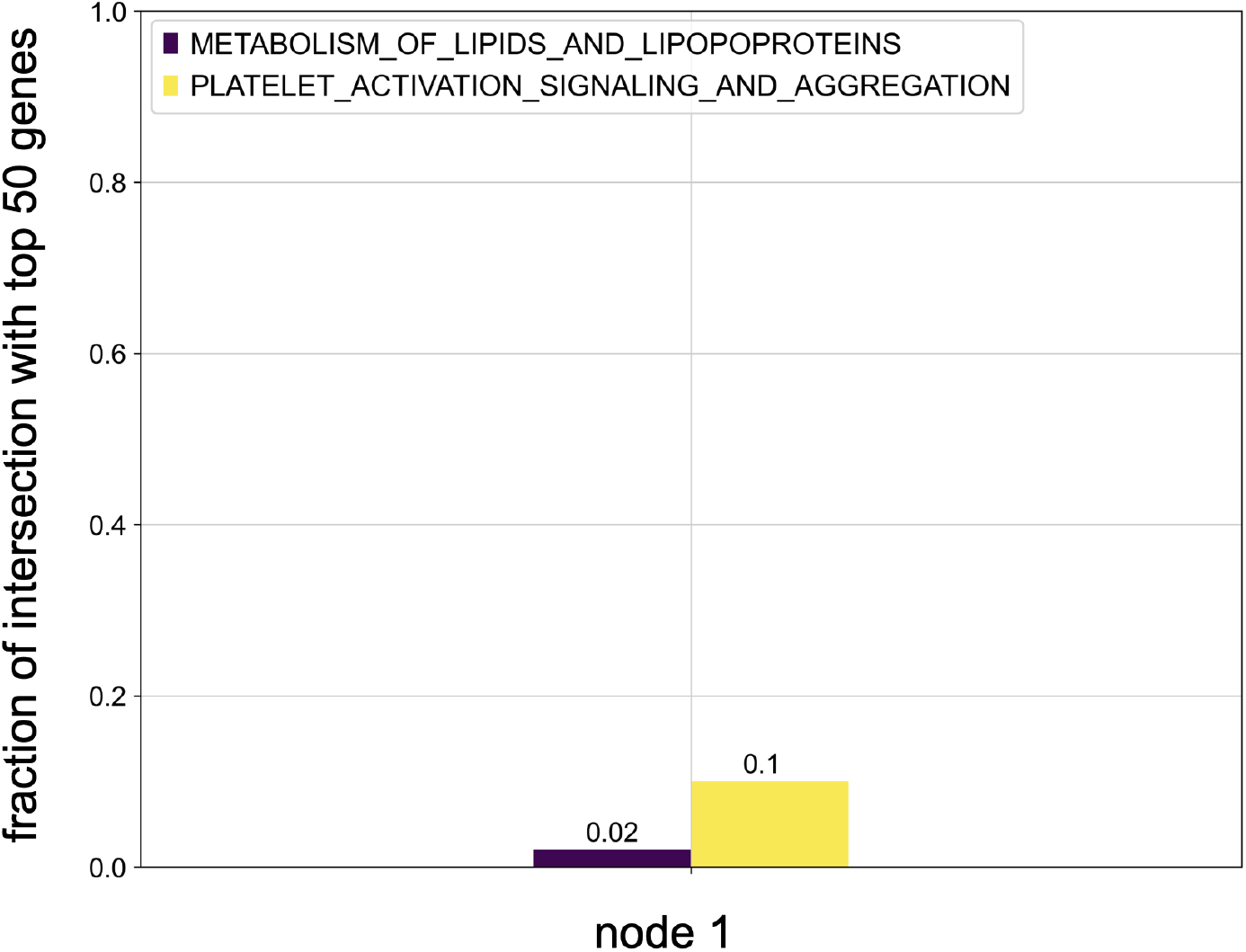
Learning new GPs for query data. Comparison of the top influential genes dominating the variance in node 1 with genes from the top GPs for CD14^+^/16^+^ monocytes from **Fig. 2d**.

**Supplementary Figure 10.**
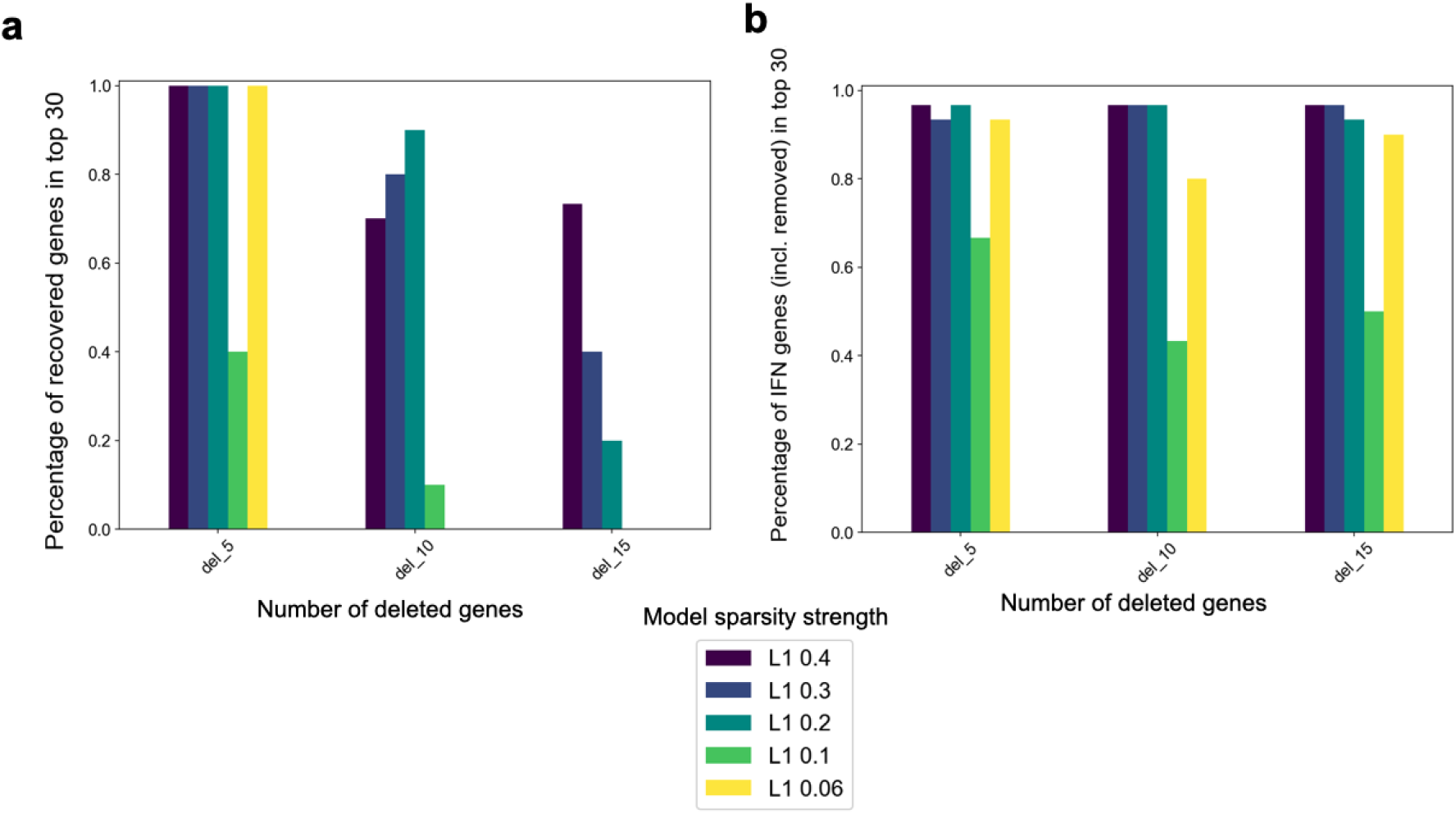
Benchmarking expiMap for enriching predefined GPs. The expiMap model was trained on the PBMCs dataset from Kang et al. (n = 13,576), with [CYTOKINE_SIGNALING_IN_IMMUNE_S’, ‘INTERFERON_ALPHA_BETA_SIGNALING’, ‘ANTIVIRAL_MECHANISM_BY_IFN_STI’, ‘INTERFERON_GAMMA_SIGNALING’, ‘IMMUNE_SYSTEM’] removed from GPs obtained from the Reactome database and only ‘INTERFERON_SIGNALING’ was kept for training. **(a)**The x-axis shows the number of deleted top genes in ‘INTERFERON_SIGNALING’ program, while the y-axis shows the percentage of those genes added to the top 30 genes in the ‘INTERFERON_SIGNALING’ program after training. The colors show the different values of L1 sparsity for each experiment. **(b)**The x-axis is the same as in **(a);**the y-axis demonstrates the percentage of the original interferon-related genes among the top 30 genes in the ‘INTERFERON_SIGNALING’ program after training. When the y-axis value is smaller than 1.0 it means that a 1–y percentage of false-positive genes was added to the GP.

**Supplementary Figure 11.**
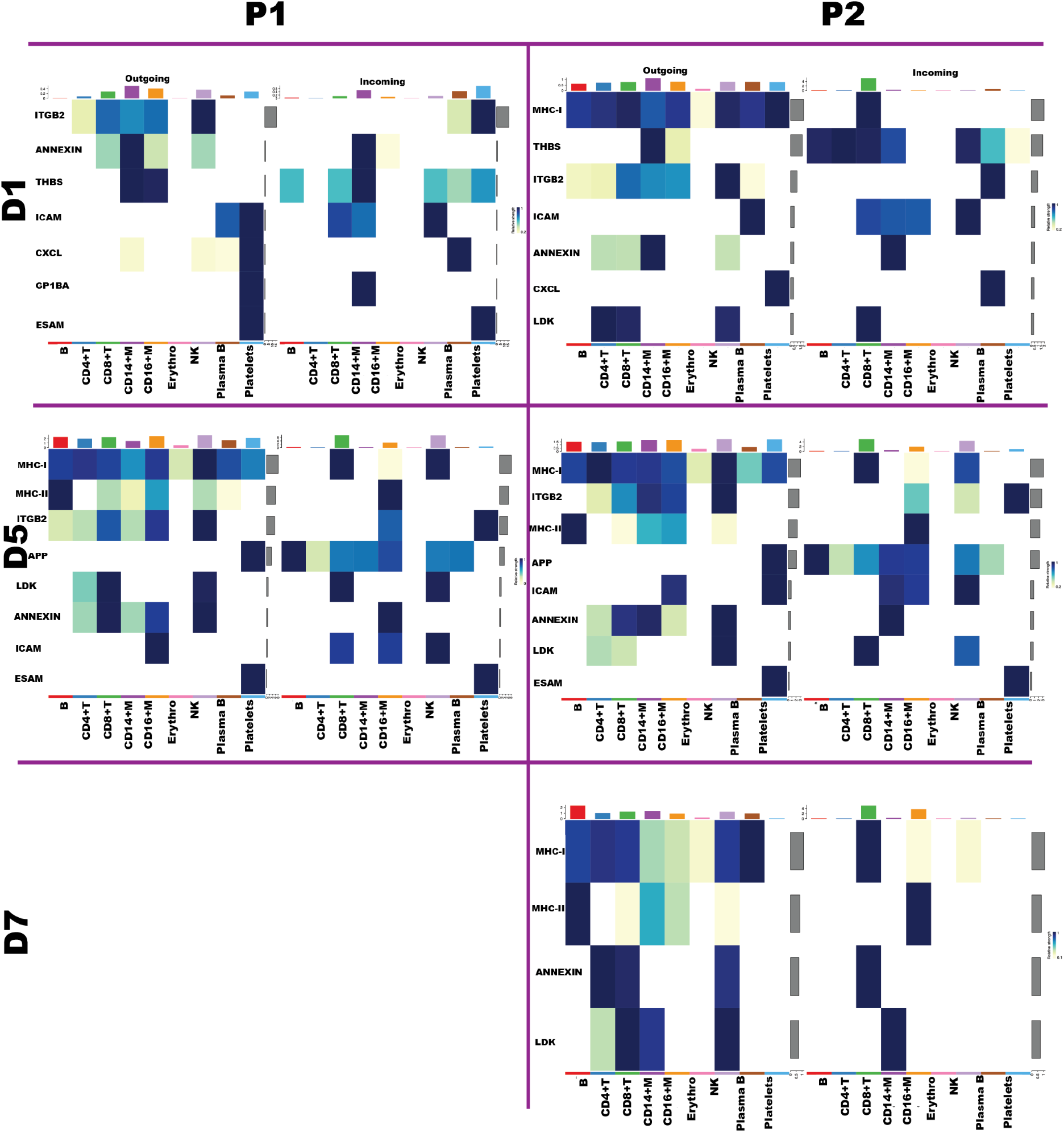
Cell–cell communication pathways in PBMCs from two patients with COVID. The heat maps show the activated cell communication circuits in two patients treated with tocilizumab during the course of admission into the ICU with COVID infection. Each column provides information for each patient and each row provides data for each day of treatment. B = B cells, CD4^+^ T = CD4^+^ T cells, CD8^+^ T = CD8^+^ T cells, CD14^+^ M = CD14^+^ monocytes, CD16^+^ M = CD16^+^ monocytes, Erythro = Erythrocytes, Plasma B = Plasma B cells, Platelets.

**Supplementary Figure 12.**
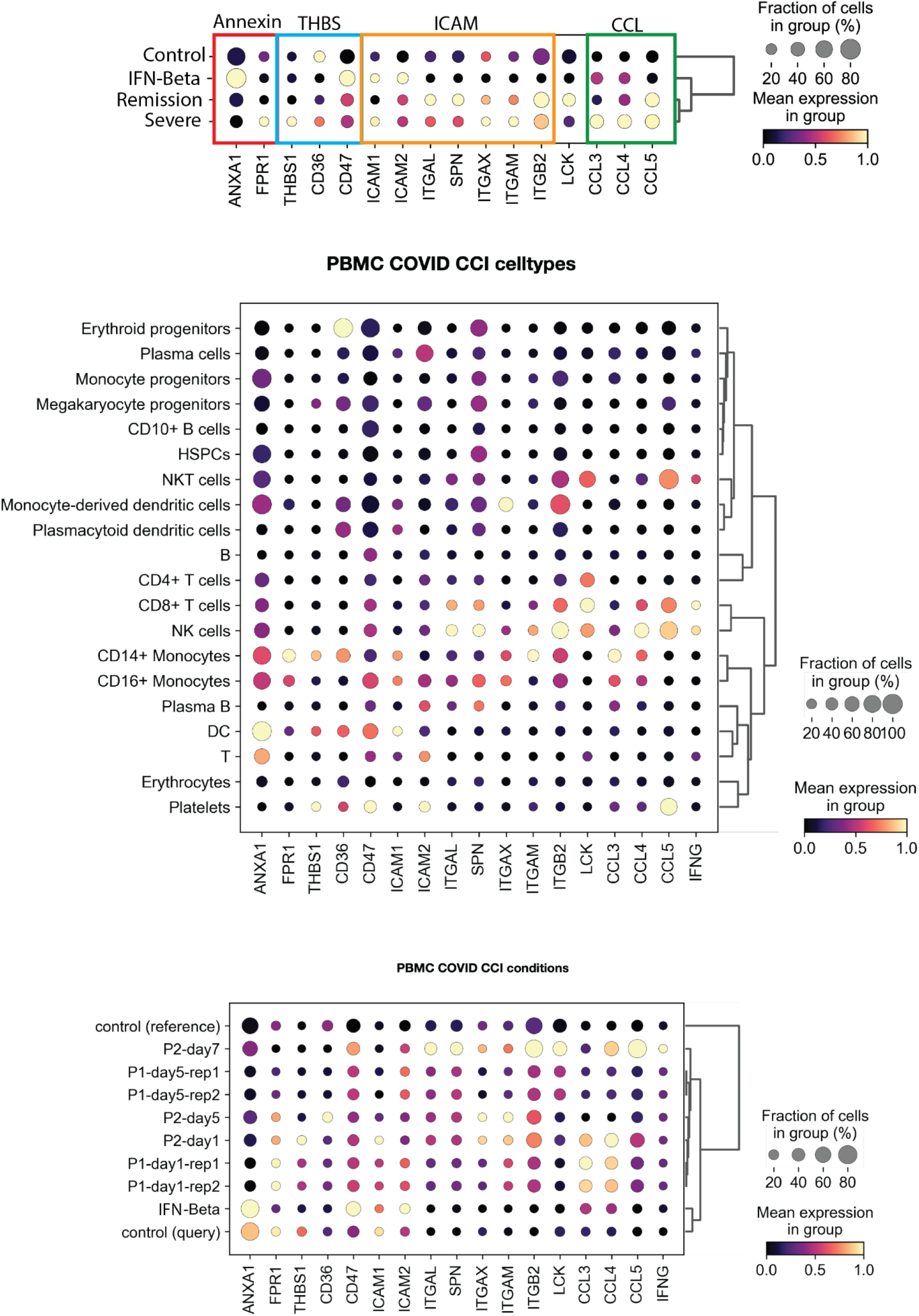
Transcriptional activity of cellular communication circuits occurring in severe COVID. Transcriptional activity of ligand–receptor pairs associated with cell–cell communication pathways and interferon gamma *(IFNG)* by annotated cell type and studied conditions. The pathways represented are annexins (*ANXA1, FPR1),* THBS *(THBS1, CD36, CD47),* ICAM *(ICAM1, ICAM2, ITGAL, SPN, ITGAX, ITGAM, ITGB2),* LCK, and CCL *(CCL3, CCL4, CCL5)*.

**Supplementary Figure 13.**
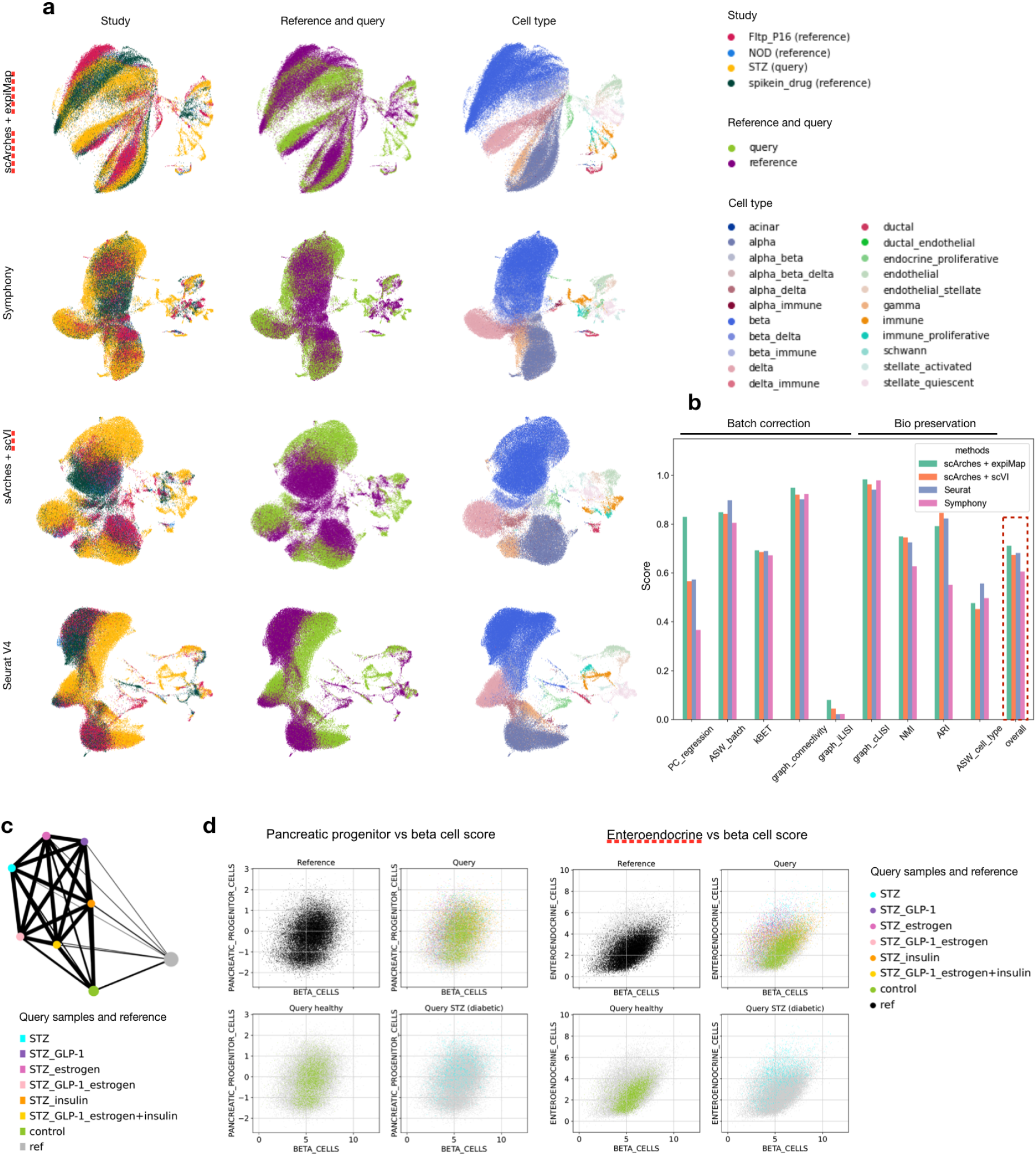
Comparison of integration results across different integration methods. **(a)** UMAPs of integrated embeddings obtained with different integration methods, **(b)**comparison of integration quality across methods, **(c)** PAGA of integrated beta cells indicates that the connection of reference cells with query control cells is the strongest, the connection of T2D-model query cells treated with insulin is moderate, and the connection with other T2D-model cells is the weakest. **(d)** expiMap term scores in beta cells correspond to the known loss of beta cell identity, dedifferentiation, and transdifferentiation in diabetes. Left, loss of beta cell identity (x-axis) vs dedifferentiation-related (y-axis) expiMap terms; right, loss of beta cell identity (x-axis) vs transdifferentiation-related (y-axis) expiMap terms.

**Supplementary Figure 14.**
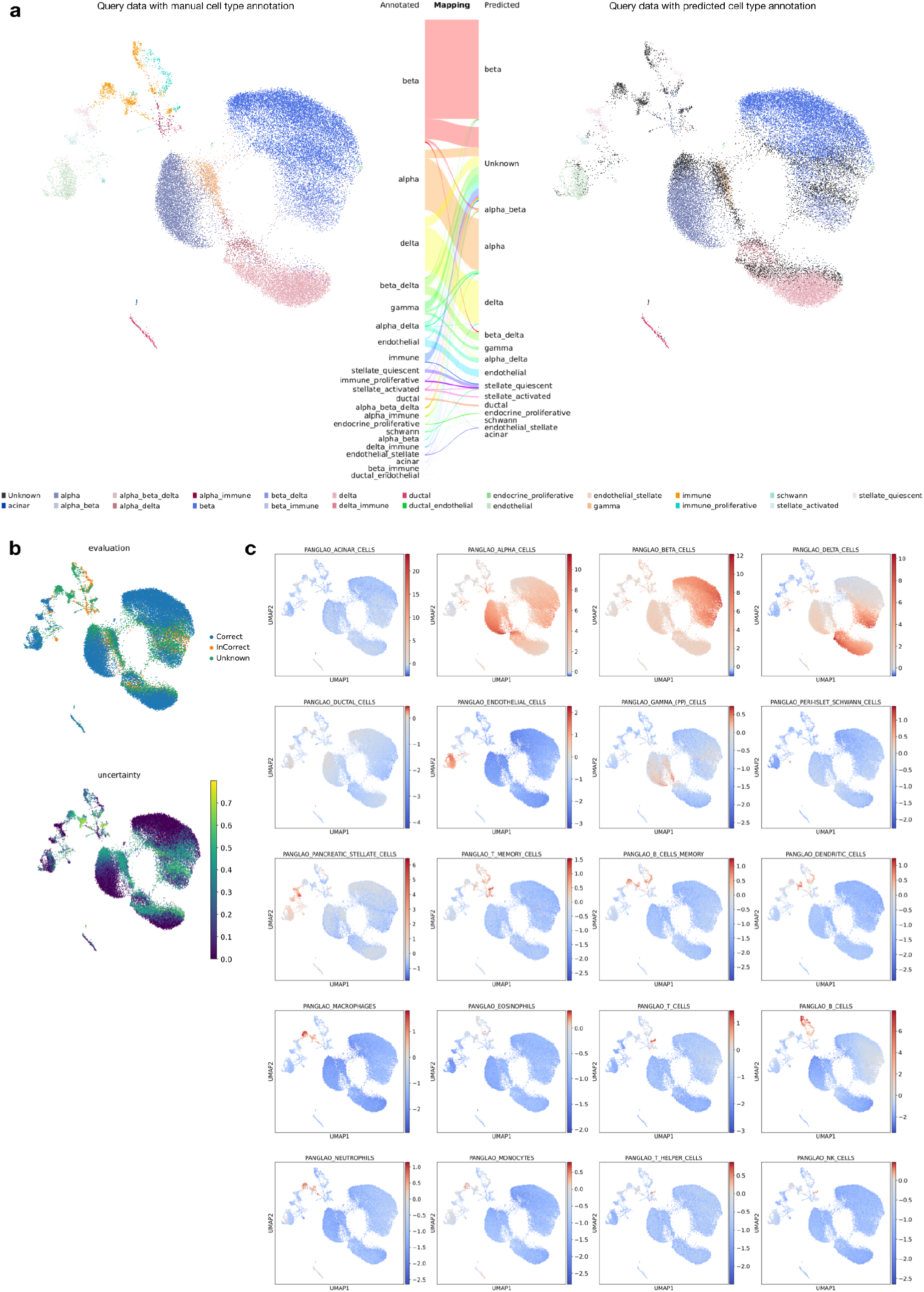
Cell type annotation based on expiMap with automatic annotation transfer and the use of cell type-specific gene set scores. **(a)** Correspondence between manual and transferred annotations in query. UMAPs were calculated using expiMap terms from PanglaoDB. **(b)** Annotation transfer success and uncertainty in query. **(c)** expiMap scores of cell types known to be present in the pancreas can be used for manual cell type annotation.

**Supplementary Figure 15.**
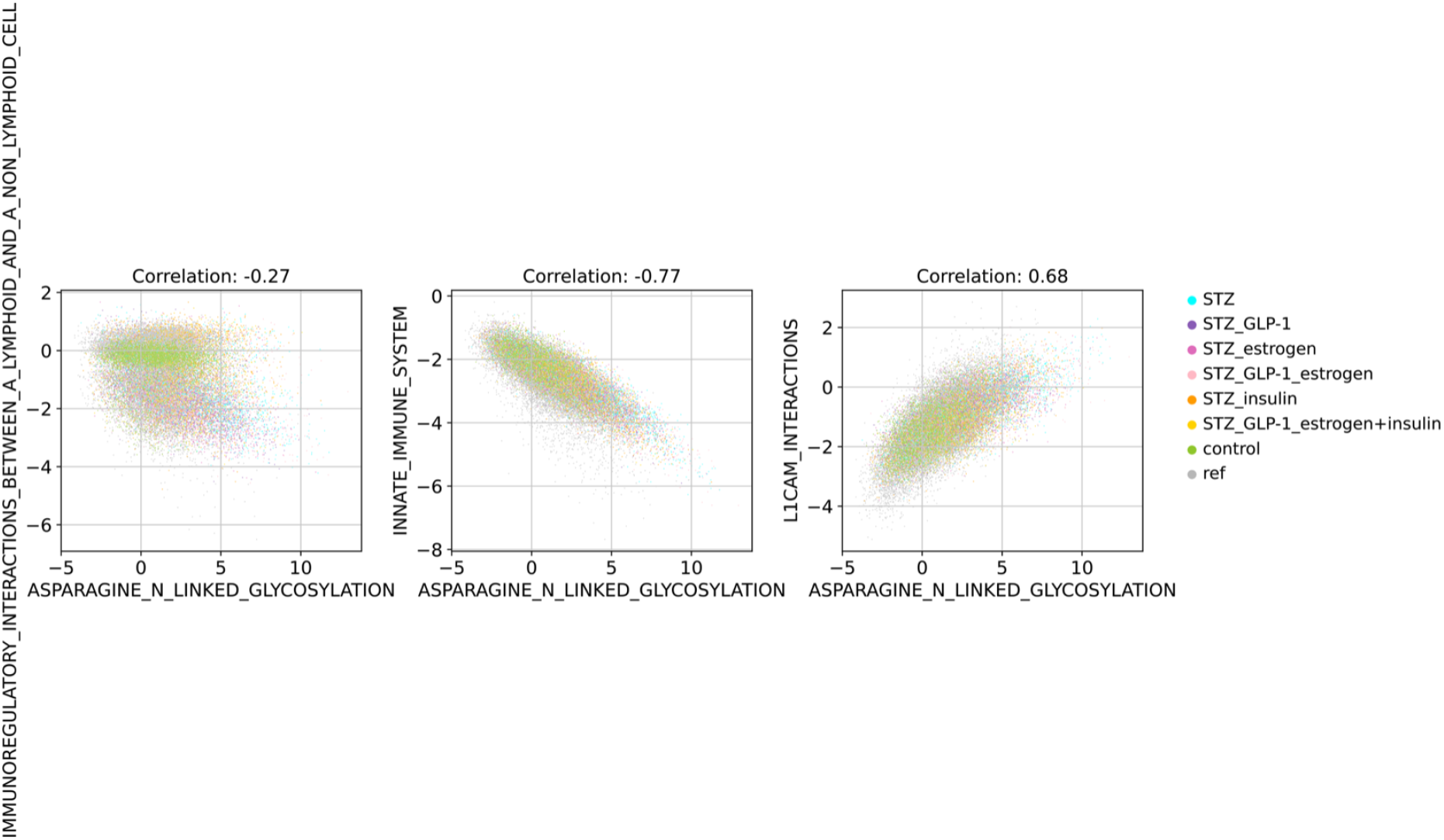
Comparison of beta cell scores of immune-related terms differentially active in T2D-model beta cells and asparagine *N*-linked glycosylation term. Ref: reference datasets, other samples are from the query dataset; STZ: streptozotocin T2D-model; STZ_GLP-1: streptozotocin T2D-model treated with GLP-1; STZ_estrogen: streptozotocin T2D-model treated with estrogen; STZ_GLP-1_estrogen: streptozotocin T2D-model treated with GLP-1-estrogen conjugate; STZ_insulin: streptozotocin T2D-model treated with insulin; STZ_GLP-1_estrogen+insulin: streptozotocin T2D-model treated with GLP-1-estrogen conjugate and insulin; control: healthy control.

**Supplementary Figure 16.**
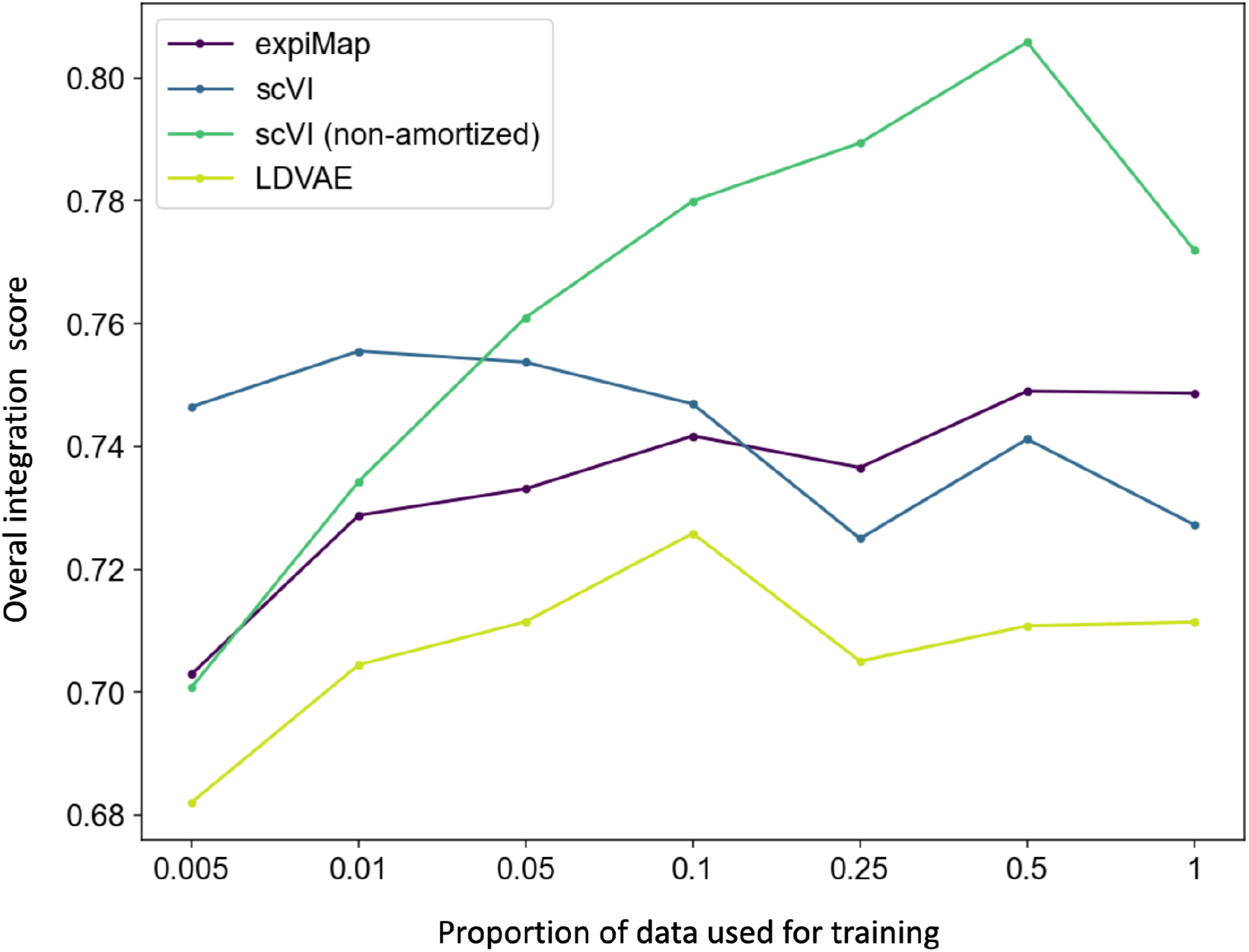
Subsampling effect on data integration. The overall integration accuracy for different subsamples of PBMCs (n = 161,764)^8^ data across different models. The x-axis denotes the proportion of the data used for training each model; the y-axis is the overall average score across nine integration metrics measuring both biological preservation and batch removal, as introduced in **Fig. 3b**.

## Expimap - Online Methods

### expiMap model

Our model builds upon the framework of (conditional) variational autoencoders [1, 2]. The log-likelihood of the data for expiMap can be written as

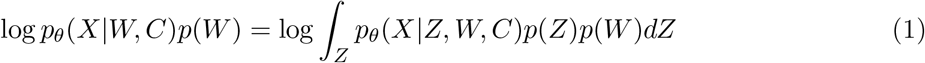

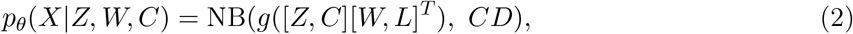

where *g*(*x*) = softmax(*x*) * *S* is a softmax function that is multiplied by the library size *S* of each cell. Alternatively, *g*(*x*) could also be a softplus or exponential function. Further, *X* is a random variable representing gene expression, *C* indicates conditions (e.g., batch id), and *p*_*θ*_(*X | Z, W, C*) is the output distribution, also called a decoder in the setting of variational autoencoders, used to model *X* given the latent variable *Z*

NB(.,.) in 2 denotes the mean and dispersion parametrized negative binomial distribution, [.,.] means a column stacked matrix, *W, L* are matrix parameters for latent variables *Z* and one-hot encoded conditions *C*, respectively; and *D* is a matrix of condition-specific dispersion parameters for each gene. *W* is a *n × m* matrix with *n* corresponding to the number of genes and *m* corresponding to the number of gene programs (GPs) provided as an input.

The prior *p*(*W*) in 1 is defined as:

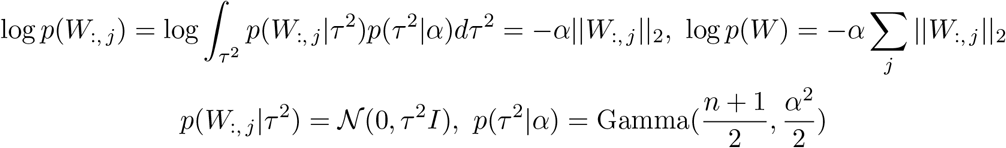

The constants were omitted because they do not affect the optimization. We use a hierarchical Bayesian prior on the columns of *W* with the parameter *τ*^2^ integrated out as in oi-VAE [3], resulting in the lasso regularization term. The lasso regularization allows the model to deactivate the GPs that do not contribute to the reconstruction loss in the model. *α* is a hyperparameter specifying the strength of the group lasso regularization.

The evidence lower bound (ELBO) is a part of our total loss to train the model. During the model training, the posterior distribution *p*_*θ*_(*Z*|*X, C*) is approximated by the variational distribution *q*_*ϕ*_(*Z*|*X, C*), which includes a deep neural network parameterized with *ϕ*; it is also called an encoder.

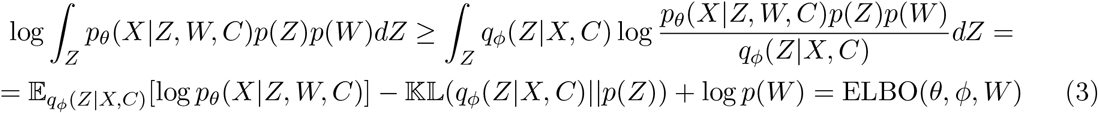

Where *θ, ϕ* are parameters of the decoder and the encoder, respectively.

### Gene program matrix

We use tab-delimited text files w here the r ows r epresent g ene s ets a s a n i nput to c onstruct masks for *W* (see the previous section). The first column is reserved for the name of the gene sets and the other columns should contain the names of genes. GMT files could be directly used in our api as an input.

A database could be also passed to the model in the form of a binary matrix *B* with columns corresponding to GPs and rows corresponding to genes, with *B*_*i,j*_ = 1 if the *i*th gene is in the *j*th gene program and 0 otherwise. Such a matrix is actually always constructed from the files described above before passing to the model. We refer to matrix *B* as the GP matrix.

### Defining hard/soft gene membership

The decoder network in 2 consists of a linear layer *H* = [*Z, C*][*W, L*]^*T*^, in which the output is then transformed to a negative binomial means by the nonlinear function *g*(*H*). The GP matrix *B* specifies gene programs and the gene memberships for these programs. The matrix *B* is used as a mask for the matrix of the decoder weights *W*, where the parameters for inactive genes in each gene program are set to zero and do not change during training if the hard mask is used.

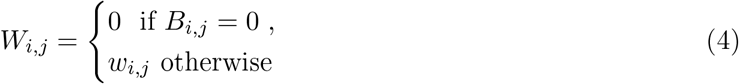

In the case of a soft mask, we add a regularization term that forces gene weights for genes that are not originally part of a GP to become zero, but also allows them to become active (non zero) if they contribute to the reconstruction:

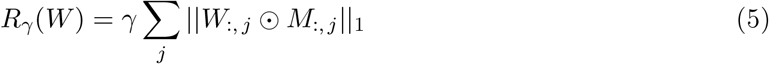

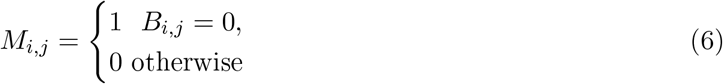

Some columns of M can be set to a vector of ones 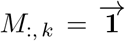 by setting 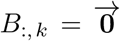 to allow the introduction of sparse GPs.

Both variants (hard and soft masks) force the columns of *Z* to correspond to the GPs encoded in *W*.

### Learning new gene programs

To allow new GPs to be learned, the model can be extended with additional nodes in reference training or query projection. For this, the last layer of the encoder is expanded with additional nodes connected to the existing nodes from the previous layer and producing the new vector *Z*_new_; in the decoder, the additional matrix *W*_new_ is concatenated to *W* (now denoted by *W*_old_), resulting in:

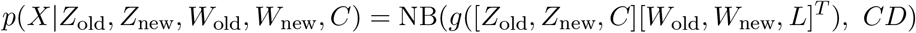

In addition, L1 regularization is added to *W*_new_, which is equivalent to the Laplace prior to this matrix. In addition, for each element of the vector *Z*_new_ the sample estimate of Hilbert Schmidt independence criterion (HSIC) between the element and the other elements of *Z*_old_ and *Z*_new_ is added as a regularization term to the loss [4]. Also *W*_new_ can be constrained with hard gene membership or regularized with soft gene membership (see the previous section) as *W*_old_ using an additional gene program database. In this case we don’t use HSIC regularization for these new constrained nodes.

### Training

We use the stochastic proximal gradient descent to optimize the ELBO loss 3 with additional regularization terms. We also multiply the KL divergence in the ELBO loss by the regularization coefficient *β*. Excluding the group lasso *R*_*α*_(*W*) = − log *p*(*W*) and soft mask term *R_γ_* (*W*) that appear in the proximal update step (discussed further), the loss function of the model can be written as:

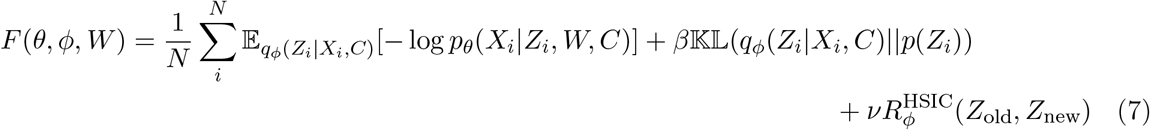

Where 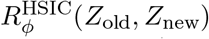 is a sample estimate of the HSIC regularization term. In addition, *Z*_new_ and *Z*_old_ are the old (existing in reference model) and new (learned in query training) unconstrained programs, respectively.

Then, to minimize the objective function we use the update scheme

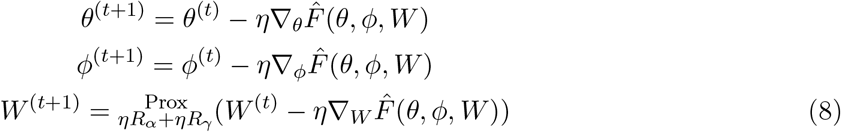

Where 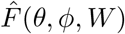 denotes an estimate of the function 7 over a mini-batch of samples (as in the standard stochastic gradient descent algorithm), and *η* is a learning rate.

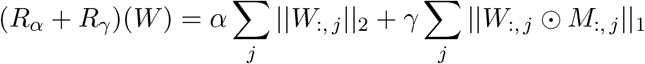

is the lasso and soft mask regularization term and its proximal operator is

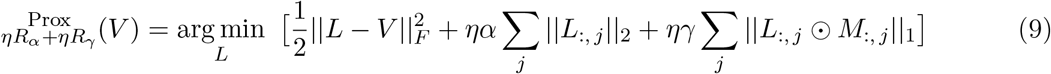

The hard mask variant implies *γ* = 0. The proximal operator above has a closed-form expression (see the next section for the derivation), so it is easy to apply it after the stochastic gradient descent update. The gradient for the expectation terms is obtained with the reparametrization trick, as is common in the VAE framework[1].

### Proximal operators for expiMap

To derive the closed form of the proximal operator described in the previous section, we need two theorems.

#### Theorem 1 (Proximal operator of separable functions)

*Suppose that f*: *E*_1_ × *E*_2_ ×…× *E*_*m*_ → (−∞, ∞] *is given by*

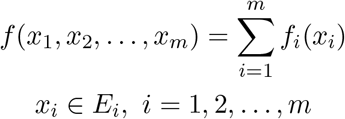

*Then for any x*_1_ ∈ *E*_1_, *x*_2_ ∈ *E*_2_,…, *x*_*m*_ ∈ *E*_*m*_,

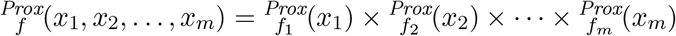

where *E*_*i*_ denotes a vector space, and × is a Cartesian product. The proof of this theorem can be found in [5].

#### Theorem 2 (Decomposition of the proximal operator)

*A sufficient condition for* 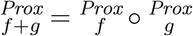

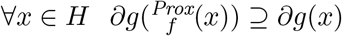

Where *f, g* are closed (or, equivalently here, continuous), convex functions; *H* denotes a Hilbert space, and *∂g* stands for a subgradient of *g*.

The proof of the theorem can be found in [6].

We use the two theorems above to find the closed form of the proximal operator 9. The explicit form of the regularization function is:

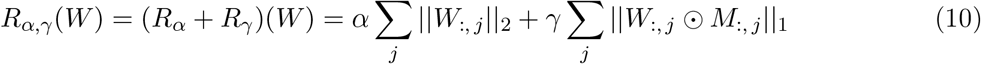

The sums in the regularization function are made over columns of *W*; thus, this function is clearly separable in columns, and theorem 1 is applicable here. This means that we only need to calculate the proximal operator for a column, as we can find the full proximal operator as a Cartesian product of the proximal operators for different columns. This is the same as using its own proximal operator for each column of *W* separately.

The regularization summand for a separate column *k* of *W* can be written as

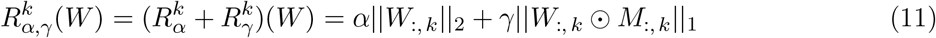

The regularization summand 11 has the form of a sum, so theorem 2 has to be used.

For the group lasso part *α*|| . ||_2_ the proximal operator can be immediately obtained (from [5]) as

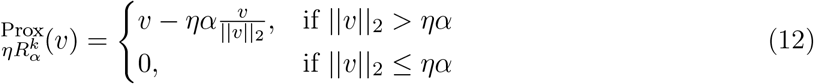

It should be noted that in the case when the mask’s column equals a vector of ones 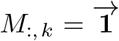 the proximal operator for the second summand in 11 *γ*| . |_1_ is just a proximal operator for a standard L1 regularization and can be written (from [5]) as

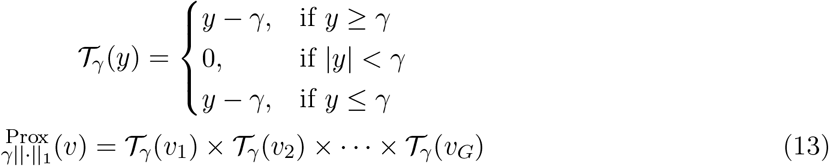

In addition, the subgradient *∂*(*γ*∨*v*∨_1_) (from [5], rewritten) is

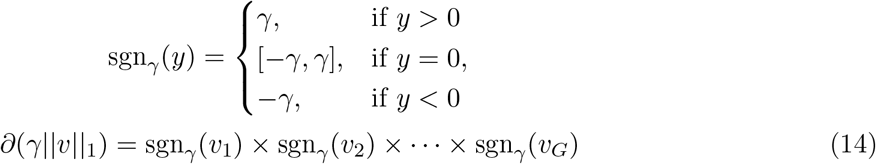

The proximal operator 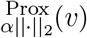 is equal to 12 (without *η*). By direct calculation for 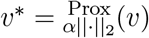 the following holds ∀*i* = 1,…, *G*: if *v*_*i*_ = 0, then 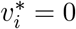 if *v*_*i*_ < 0, then 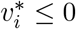; if *v*_*i*_ > 0, then 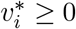.

This basically means that 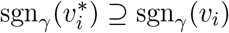. It immediately follows from the form of the subgra-dient 14 that

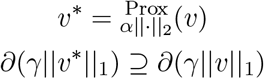

Using this and theorem 2 we can conclude that

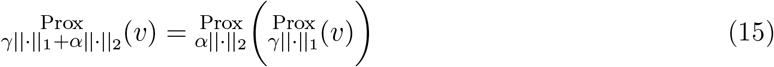

Therefore, the closed form of the proximal operator for the case *M*_:, *k*_ = **1** is:

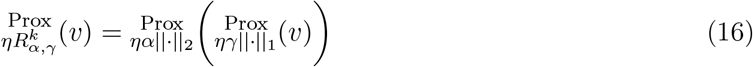

Moreover, the closed forms of 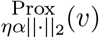 and 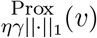 are given in 12 and 13, respectively.

For the case 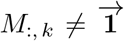, similar reasoning can be applied. First, the closed form of the proximal operator 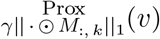 for L1 norm of the vector of genes (gene weights in the factor) that are inactive in the annotation for the factor *k* can be written as

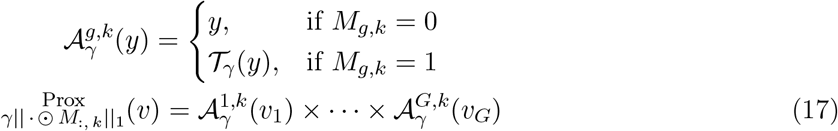

Where 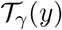 is the same as in13.

The subgradient *∂*(*γ*∥ *v* ⊙ *M*_:, *k*_∥_1_) can be written as

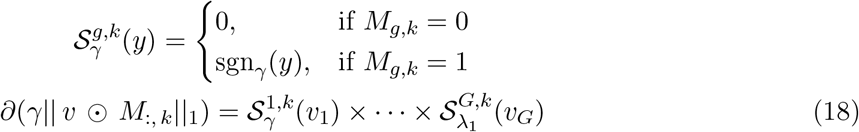

Where sgn_*γ*_(*y*) is the same as in 14.

Using the same reasoning as in the derivation of the proximal operator for sparse unannotated factors, we see that

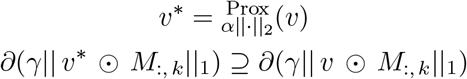

This means that we again can use theorem 2 and obtain the closed form of the proximal operator (with the learning rate *η*) for the column *k* of *W*, which corresponds to the annotated factor *k*

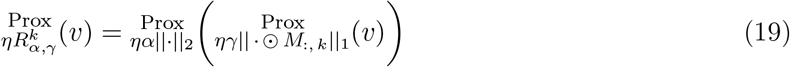

In addition, the closed forms of 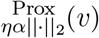 and 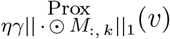 are given in 12 and 17 respectively.

Theorem 1 allows calculation of the output of the joint proximal operator 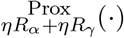 in 8 by applying the proximal operators 12, 16 or 19 on each column of the input of the joint operator independently.

### Reference mapping

The projection of a query dataset to a reference dataset is performed using the single-cell architectural surgery (scArches) approach [7]. After training a conditional VAE model for multiple batches of the reference dataset, the trained weights are transferred to a new model with additional conditional nodes used to map new query batches to the reference. Further, additional nodes for new learnable GPs can be added at this stage (see the learning new gene program section). During the training of the expanded model for query projection, only the conditional weights connecting new batches and the weights for new GPs (if any) in both encoder and decoder are tuned; the rest of the weights are frozen. Projecting with scArches preserves the latent representation of the reference and projects the query data to the same latent space while correcting for batch effects between the query and data.

### Differential testing for gene programs

To test the hypothesis *H*_0_: *Z*_*i,a*_ > *Z*_*i,b*_ versus *H*_1_: *Z*_*i,a*_ ≤ *Z*_*i,b*_, where *Z*_*i,a*_, *Z*_*i,b*_ are the dimension *i* of the latent variables for the cells from the groups *a* and *b* respectively, we use the logarithm of the Bayes factor:

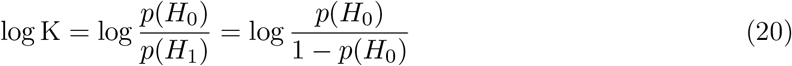

where *p*(*H*_0_) and *p*(*H*_1_) are the probabilities of the hypotheses *H*_0_ and *H*_1_ respectively. We can compute *P* (*H*_0_) as:

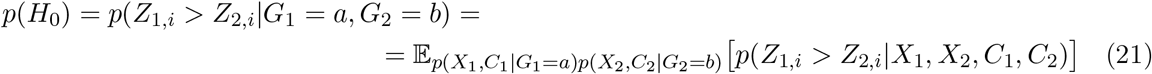

where *G*_1_ and *G*_2_ denote the independent group variables for *X*_1_ and *X*_2_, respectively.

The probability *p*(*Z*_1,*i*_ > *Z*_2,*i*_|*X*_1_*, X*_2_, *C*_1_, *C*_2_) inside the expectation in 21 can be estimated with the approximate posteriors *q*_*ϕ*_(*Z*_1,*i*_|*X, C*) and *q*_*ϕ*_(*Z*_2,*i*_|*X, C*), as follows:

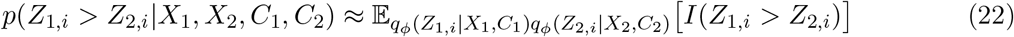

where the expectation could be approximated by sampling or calculated from the closed-form. When *q*_*ϕ*_(*Z*_*i*_|*X, C*) is Gaussian, we can calculate the expectation by

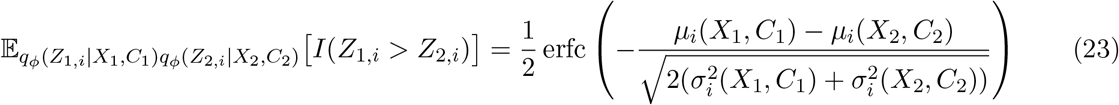

The probabilities 22 can be averaged over many cells from both groups as in scVI [8], to obtain the approximate value for 21.

Through the examples in the paper, we refer to the results obtained at the threshold log K ≥ 2.3 as the “expiMap test results”, and call such gene programs “differential GPs” in the comparison of interest in this work.

### Gene importance score

Each column of the weight matrix *W* in the decoder 2 corresponds to a gene program and each row corresponds to a gene. Because of the linearity of the decoder, a change in the latent score of the *i*th gene program *Z*_*i*_ affects the reconstruction of gene counts more for those genes with higher absolute values of the weights in *W*_:,*i*_. Consequently, we can rank genes in each gene program by the absolute values of their weights in *W*. This ranking reflects the relative importance of a given gene program for each gene; a higher ranking means that this gene is affected more by the gene program.

### Latent scores directions

The signs of latent scores of GPs do not necessary correspond to up- or downregulation of these GPs. However, in some cases, it is possible to determine if an increase in a latent score corresponds primarily to an increase or decrease in the expression of genes of a corresponding gene program. This can be determined by analyzing the decoder gene weights in the column corresponding to the gene program (as described in the previous section). If most of the gene weights in the column of *W* corresponding to the gene program are positive then the higher positive latent score implies upregulation; in the opposite case of mostly negative weights, a lower negative score also means upregulation.

For the *j*th gene program, the direction *D*_*j*_ of predominant upregulation (negative or positive) can be calculated heuristically by several methods, we use two methods:

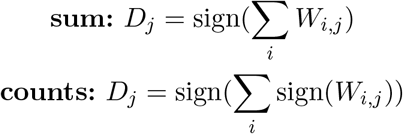

Then, we can multiply the latent score of the gene program by this direction 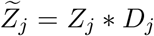, so that a higher positive value of 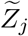 always corresponds to predominant upregulation of the gene program and a lower negative value to downregulation. These normalized scores can then be used for plotting or testing.

### Integration evaluation

Integrations were evaluated with methods implemented in scIB. We evaluated biological conservation through graph cLISI, normalized mutual information (NMI), adjusted Rand index (ARI), and average silhouette width (ASW) for cell type; and batch correction through principal component regression, average silhouette width (ASW) for batch, kBET, graph connectivity, and graph iLISI. All metrics are further described in the scIB paper [9]. The overall score was computed as the average of all scores.

### Non-amortized scVI

We compared the integration performance of expiMap with scVI and non-amortized scVI. Non-amortized scVI is a VAE model similar to scVI, where the neural network encoder was replaced by a per cell vector of parameters for each cell in a dataset.

For each cell *i* there are vectors 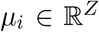 and 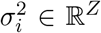 with the size of the latent space. The *j*th latent variable for the cell *i* is obtained by sampling independently from the Gaussian distribution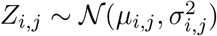. The decoder is the same as in the standard scVI model. The non-amortized scVI model is trained by minimizing the negative ELBO in batches with a gradient descent algorithm as a standard VAE model.

### Gene set enrichment analysis using limma-fry

Read counts were normalized using the trimmed mean of M-values (TMM) [10] with singleton pairing implemented in edgeR [11] to account for sparsity in the single cell RNAseq data. The fry test (Fast Approximation to ROAST) [12] in limma [13] R/Bioconductor package was applied to log counts-per-million (logCPM) values obtained by voom transformation [14] to test for the enrichment of the gene set terms in the Reactome pathway databse [15]. The Reactome database was obtained from the Molecular Signature Database (MSigDB) [16, 17].

### Supplementary note 1: comparison with limma-fry

We compared the Bayes factors from the expiMap model to the enrichment results (FDR values) from limma-fry [12, 13] applied to the PBMC IFN-*β* data using identical terms selected from Reactome.

We considered the following comparisons in the PBMC IFN-*β* data for the gene set enrichment analysis: terms enriched at the global level in IFN-*β* stimulated cells compared to control unstimulated cells; terms enriched in B cells, CD14+ Monocytes and CD16+ Monocytes where each of these populations was separately compared to all other cell-types; and terms enriched in CD14+ Monocyte population in IFN-*β* stimulated cells compared with the control (unstimulated) cells. In each of these comparisons, we added “study” as a covariate in the design matrix of the linear model in the limma-fry framework. The threshold for the absolute natural logarithm of the Bayes scores from expiMap was set to 2.3, which corresponds to a strong evidence for the enrichment of a term in one group of cells compared to another group of cells in a Bayesian hypothesis testing framework. We refer to the results obtained at thresholds larger than this nominated threshold as the “expiMap test results”, and shall call such gene programs “differential GPs” in the comparison of interest in this work. The threshold criteria for the limma-fry results was set to a mixed FDR (that is, direction-independent enrichment) value of 0.05. We chose to perform GSEA using the limma-fry pipeline to account for the complex experimental design of the integrated data, which would not otherwise have been possible with rank-based enrichment tests.

We observed that the enrichment results generally agree for both expiMap and the limma-fry framework for GSEA. However, expiMap tends to select differential GPs that are more specialsion compared to GPs enriched by fry-based GSEA, where the increased variance of gene expression measurements on the integrated atlas due to biological and technical variability can impede the detection of relevant biological signals. In **Supplementary Fig. 4a**, where IFN-*β* treated cells are compared with control (untreated) cells, the terms INTERFERON_SIGNALING and IN-TERFERON_ALPHA_BETA_SIGNALING were found to be enriched in IFN-*β*stimulated cells in both the expiMap and GSEA results. The limma-fry framework for GSEA identified the enrichment of CYTOKINE_SIGNALING_IN_IMMUNE_SYSTEM and IMMUNE_SYSTEM in IFN-*β* cells, which are considered broader and less specialized terms than the ones detected by expiMap. The terms enriched in stimulated CD14+ Monocytes were largely similar in the expiMap test and fry results (**Supplementary Fig. 4b**); however, expiMap detected the important pathway METABOLISM_OF_CARBOHYDRATES, which was missed in the fry results. We also observed that fry tends to assign significant scores to the general GPs with larger numbers of genes. For B cells and CD16+ Monocytes (**Supplementary Fig. 4c, d**), fry detects only the general terms, such as ADAPTIVE_IMMUNE_SYSTEM and IMMUNE_SYSTEM (and additionally HEMOSTASIS for CD16+ Monocytes), whereas the expiMap test identified smaller-sized and more specialized GPs for both cell types. For example, for B cells, the expiMap test identifies enriched terms such as SIGNALING_BY_B_CELL_RECEPTOR_BCR and MHC_CLASS_II_ANTIGEN_PRESENTATION (**Supplementary Fig. 4c**). The enriched Reactome GPs were almost identical in the cell type specific test for CD14+ Monocytes (**Supplementary Fig. 4e**) in the expiMap and fry approaches. In addition to an enhanced capability for detection of specialized cellular and molecular programs, expiMap removes the need to repeat differential gene expression and gene set enrichment testing for every single comparison, thereby resulting in a shorter computational time and faster data analysis.

### Datasets and preprocessing

All the cell type labels and metadata were obtained from original publications unless specifically stated below.

### Immune healthy atlas

The immune dataset includes samples from bone marrow cells and peripheral blood cells from different human samples. The bone marrow data were collected from Oetjen et al. [18] and PBMC samples were obtained from 10x Genomics https://support.10xgenomics.com/single-cell-gene-expression/datasets/3.0.0/pbmc_10k_v3, Freytag et al. [19], and Sun et al. [20]. The detail of the retrieval path and the preprocessing can be found in Luecken et al. [9] and Lotfollahi et al. [7] We used the Reactome pathway database for annotations [15] from MsigDB [16, 17]; we also removed all pathways with fewer than 12 genes. The genes that were not in the GPs database were filtered out, reducing the total number of genes from approximately 11,000 to 3690. Then 2000 highly variable genes were selected for training.

### PBMC IFN-*β*

This dataset contains cells from eight patients with Lupus treated with IFN-*β* or left untreated for six hours [21]. The pools from the IFN-*β* and control cells were mixed together and loaded to a 10x kit. The dataset was obtained from the Seurat tutorial (https://satijalab.org/seurat/articles/integration_introduction.html). We have used the same genes as in the reference (Immune atlas).

### PBMC COVID-19

The dataset [22] contains five peripheral blood samples from two patients with severe COVID-19 at three different time points, consisting of severe remission during treatment with tocilizumab. The blood samples were collected on day 1, within 12 h of tocilizumab treatment, and on day 5 for both patients. An additional blood sample was collected from patient 2 because the patient remained COVID-19 positive. The cell types were annotated using markers provided by the authors in the original study. We have used the same genes as in the reference (Immune atlas). The dataset is available on GEO; the accession number is GSE150861.

We used the integrated dataset to analyze cell-cell interactions using the CellChat package [23]. For this analysis, we used the non-integrated shared gene space between all the integrated datasets after removing those genes supported by less than five counts, for a total of 10,851 genes ready for analysis. We then ran CellChat on each subset using the curated database for interactions in human samples. The gene expression of each ligand-receptor pair was visualized using a dotplot generated using Scanpy 1.8.1 [24] and anndata 0.7.6 [25]. The scripts for the analyses, as well as the package version used in the analysis, can be found in the “covid” section of the repository.

### Pancreas

The datasets are publicly available on GEO and further described in **Supplementary Table 4**. We removed low-quality cells (high mitochondrial fraction, low number of genes) using a study-specific threshold. For cell type annotation, we removed genes expressed in fewer than 20 cells in each study and normalized the expression in each study to 1e6 total counts, excluding highly expressed genes, and subsequently applied a log transformation. We merged datasets across studies using Ensembl IDs and retaining the genes expressed in all studies. We used merged data across studies, followed by the identification of highly variable genes, z-normalization, and the computation of top PCA components. We clustered the data and plotted known pancreatic islet cell type markers to annotate cell types cluster-wise.

For integration, we separated the datasets into reference and query, as described in **Supplementary Table 4**. From the reference data, we removed immune cell types and their doublets. We removed genes expressed in fewer than 20 cells in the reference data. We used gene sets from PanglaoDB [26] release from March 2020 and Reactome [15] v4.0, and mapped them to mouse genes using Ensembl [27] V103 orthologs. We used only gene sets with at least three genes and at most 200 genes. We excluded genes that were not present in these gene sets. With expiMap, we integrated the reference datasets using samples as batches and projected query samples. We also performed matched integrations with Seurat [28], Symphony [29], and scVI [8]. We evaluated different integrations, as described in the integration evaluation section. We used reference query split as batches and excluded non-healthy query samples as they were not expected to be integrated into the healthy reference owning to biological differences. For the downstream interpretation analysis, we used directed expiMap scores.

We used multiple methods to evaluate PanglaoDB cell type scores. We plotted the PanglaoDB cell type scores of expected pancreatic cell types on query UMAPs and visually compared the results to cell type annotation. We used the PanglaoDB gene set scores as features for the annotation transfer from reference to query with weighted KNN. We evaluated the annotation transfer with F1 score and by visual evaluation of prediction accuracy and certainty on UMAP.

### Integration benchmark datasets

We leverage datasets from five different tissues including PBMCs (n=161,764) [28], heart (n=18641) [30], lung (n=65,662) [31], colon (n=34,772) [32], and liver (n=113,063) [33]. All datasets, except heart, were obtained from the Sfaira database [34], which includes cell type labels. Heart was obtained from the scVI package. For the expiMap training for each dataset we used the Reactome pathway database, selected only pathways that contain more than 12 genes and filtered out all genes that are not present in any pathway, and then we selected 2000 HVG for training. For the other models, we used the same lists of genes.

### Hyperparameters

This section describe the hyperparamters used for the training models in different scenarios.

---

**Supplementary Table 5.** The label type used for each experiment as conditions for expiMap models.

| Experiment | Condition label |
| --- | --- |
| Immune Atlas (reference) + PBMC IFN-β (query) | study |
| Immune Atlas (reference) + PBMC IFN-β & COVID (query) | study |
| Pancreas | study_sample |
| PBMC IFN-β (deleted genes recovery) | study |
| PBMCs (Integration evaluation) | donor |
| Heart (Integration evaluation) | cell_source |
| Lung (Integration evaluation) | tech_sample |
| Colon (Integration evaluation) | Source |
| Liver (Integration evaluation) | orig.ident |

**Supplementary Table 6.** Usage of features for each experiment.

| Experiment | soft mask | group lasso | new GPs |
| --- | --- | --- | --- |
| Immune Atlas (reference) + PBMC IFN-β (query) | No | Yes | No |
| Immune Atlas (reference) + PBMC IFN-β (query) | Yes (query) | Yes | Yes (3 in query) |
| Immune Atlas (reference) + PBMC IFN-β & COVID (query) | No | Yes | No |
| Pancreas | No | Yes | No |
| PBMC IFN-β (deleted genes recovery) | Yes | Yes | No |
| Integration evaluation | No | Yes | No |

**Supplementary Table 7.** expiMap detailed architecture for the Immune Atlas (reference) + PBMC IFN-*β* (query) experiment (**figure 2**).

| Name | Operation | NoF/Kernel Dim. | Dropout | LN | Activation | Input |
| --- | --- | --- | --- | --- | --- | --- |
| **Inputs** |  |  |  |  |  |  |
| data | - | #Genes | × | × | - | - |
| conditions | - | #Conditions | × | × | - | - |
| **Encoder** |  |  |  |  |  |  |
| Layer_1 | FC | 256 | 0.05 | ✓ | ReLU | [data, study labels] |
| Layer_2 | FC | 256 | 0.05 | ✓ | ReLU | Layer_1 |
| Layer_3 | FC | 256 | 0.05 | ✓ | ReLU | Layer_2 |
| mean | FC | #Gene Programs | × | × | Linear | Layer_3 |
| var | FC | #Gene Programs | × | × | Linear | Layer_3 |
| latent | Multivariate Normal | #Gene Programs | × | × | - | [mean, var] |
| **Decoder** |  |  |  |  |  |  |
| Layer_1 | FC | #Genes | × | × | softmax | [latent, study labels] |
| predicted count mean | Multiplication | #Genes | × | × | - | [Layer_1, library scale] |
| **Hyperparameters** |  |  |  |  |  |  |
| Loss | NB |  |  |  |  |  |
| Optimizer | Adam |  |  |  |  |  |
| Learning Rate | 0.001 |  |  |  |  |  |
| epsilon | 0.01 |  |  |  |  |  |
| Batch Size | 128 |  |  |  |  |  |
| # of Epochs | max. 400 |  |  |  |  |  |
| alpha (group lasso weight) | 0.7 |  |  |  |  |  |
| alpha_kl (KL term weight) | 0.5 (reference) | 0.1 (query) |  |  |  |  |
| alpha_epoch_anneal (epochs for alpha_kl annealing) | 100 |  |  |  |  |  |

**Supplementary Table 8.** expiMap detailed architecture for the Immune Atlas (reference) + PBMC IFN-*β* (query) learning new GPs experiment (**figure 4**).

| Name | Operation | NoF/Kernel Dim. | Dropout | LN | Activation | Input |
| --- | --- | --- | --- | --- | --- | --- |
| **Inputs** |  |  |  |  |  |  |
| data | - | #Genes | × | × | - | - |
| conditions | - | #Conditions | × | × | - | - |
| **Encoder** |  |  |  |  |  |  |
| Layer_1 | FC | 300 | 0.05 | ✓ | ReLU | [data, study labels] |
| Layer_2 | FC | 300 | 0.05 | ✓ | ReLU | Layer_1 |
| Layer_3 | FC | 300 | 0.05 | ✓ | ReLU | Layer_2 |
| mean | FC | #Gene Programs | × | × | Linear | Layer_3 |
| var | FC | #Gene Programs | × | × | Linear | Layer_3 |
| latent | Multivariate Normal | #Gene Programs | × | × | - | [mean, var] |
| **Decoder** |  |  |  |  |  |  |
| Layer_1 | FC | #Genes | × | × | softmax | [latent, study labels] |
| predicted count mean | Multiplication | #Genes | × | × | - | [Layer_1, library scale] |
| **Hyperparameters** |  |  |  |  |  |  |
| Loss | NB |  |  |  |  |  |
| Optimizer | Adam |  |  |  |  |  |
| Learning Rate | 0.001 |  |  |  |  |  |
| epsilon | 0.01 |  |  |  |  |  |
| Batch Size | 128 |  |  |  |  |  |
| # of Epochs | max. 200 (reference) | 150 (query) |  |  |  |  |
| alpha (group lasso weight) | 0.7 |  |  |  |  |  |
| gamma_ext (L1 regularization weight for new unconstrained nodes) | - | 0.7 (query) |  |  |  |  |
| gamma_epoch_anneal (epochs for gamma_ext annealing) | - | 50 (query) |  |  |  |  |
| alpha_kl (KL term weight) | 0.5 (reference) | 0.1 (query) |  |  |  |  |
| alpha_epoch_anneal (epochs for alpha_kl annealing) | 100 (reference) | 50 (query) |  |  |  |  |
| beta (HSIC weight for new unconstrained nodes) | - | 3 (query) |  |  |  |  |
| alpha_l1 (soft mask weight for new constrained nodes) | - | 0.9 (query) |  |  |  |  |
| 3 unconstrained and 1 constrained new GPs in query |  |  |  |  |  |  |

**Supplementary Table 9.** expiMap detailed architecture for the Immune Atlas (reference) + PBMC IFN-*β* & COVID (query) experiment (**figure 5**).

| Name | Operation | NoF/Kernel Dim. | Dropout | LN | Activation | Input |
| --- | --- | --- | --- | --- | --- | --- |
| **Inputs** |  |  |  |  |  |  |
| data | - | #Genes | × | × | - | - |
| conditions | - | #Conditions | × | × | - | - |
| **Encoder** |  |  |  |  |  |  |
| Layer_1 | FC | 256 | 0.05 | ✓ | ReLU | [data, study labels] |
| Layer_2 | FC | 256 | 0.05 | ✓ | ReLU | Layer_1 |
| Layer_3 | FC | 256 | 0.05 | ✓ | ReLU | Layer_2 |
| mean | FC | #Gene Programs | × | × | Linear | Layer_3 |
| var | FC | #Gene Programs | × | × | Linear | Layer_3 |
| latent | Multivariate Normal | #Gene Programs | × | × | - | [mean, var] |
| **Library Encoder** |  |  |  |  |  |  |
| Layer_1 | FC | 128 | 0.05 | ✓ | ReLU | [data, study labels] |
| mean | FC | 1 | × | × | Linear | Layer_1 |
| var | FC | 1 | × | × | Linear | Layer_1 |
| sizefactors | Normal | 1 | × | × | - | [mean, var] |
| **Decoder** |  |  |  |  |  |  |
| Layer_1 | FC | #Genes | × | × | softmax | [latent, study labels] |
| predicted count mean | Multiplication | #Genes | × | × | - | [Layer_1, sizefactors] |
| **Hyperparameters** |  |  |  |  |  |  |
| Loss | NB |  |  |  |  |  |
| Optimizer | Adam |  |  |  |  |  |
| Learning Rate | 0.001 |  |  |  |  |  |
| epsilon | 0.01 |  |  |  |  |  |
| Batch Size | 128 |  |  |  |  |  |
| # of Epochs | max. 400 (reference) | max. 200 (query) |  |  |  |  |
| alpha (group lasso weight) | 0.7 |  |  |  |  |  |
| alpha_kl (KL term weight) | 0.5 (reference) | 0.1 (query) |  |  |  |  |
| alpha_epoch_anneal (epochs for alpha_kl annealing) | 100 (reference) | 50 (query) |  |  |  |  |

**Supplementary Table 10.**
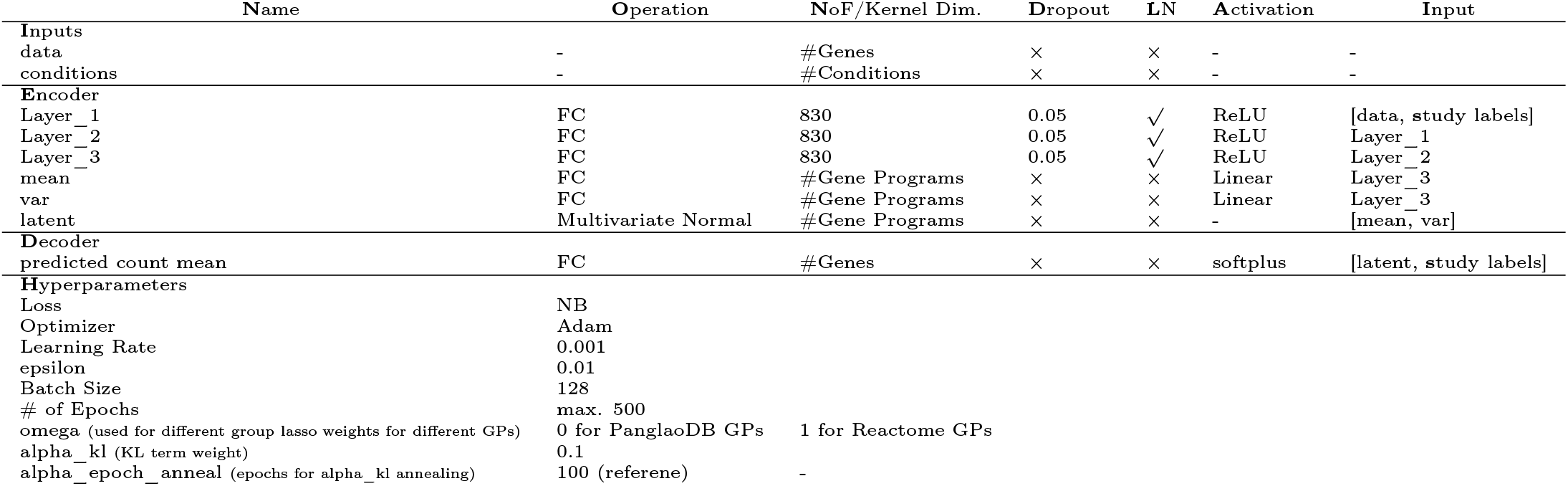
expiMap detailed architecture for the Pancreas experiment (**figure 6**).

| Name | Operation | NoF/Kernel Dim. | Dropout | LN | Activation | Input |
| --- | --- | --- | --- | --- | --- | --- |
| **Inputs** |  |  |  |  |  |  |
| data | - | #Genes | × | × | - | - |
| conditions | - | #Conditions | × | × | - | - |
| **Encoder** |  |  |  |  |  |  |
| Layer_1 | FC | 830 | 0.05 | ✓ | ReLU | [data, study labels] |
| Layer_2 | FC | 830 | 0.05 | ✓ | ReLU | Layer_1 |
| Layer_3 | FC | 830 | 0.05 | ✓ | ReLU | Layer_2 |
| mean | FC | #Gene Programs | × | × | Linear | Layer_3 |
| var | FC | #Gene Programs | × | × | Linear | Layer_3 |
| latent | Multivariate Normal | #Gene Programs | × | × | - | [mean, var] |
| **Decoder** |  |  |  |  |  |  |
| predicted count mean | FC | #Genes | × | × | softplus | [latent, study labels] |
| **Hyperparameters** |  |  |  |  |  |  |
| Loss | NB |  |  |  |  |  |
| Optimizer | Adam |  |  |  |  |  |
| Learning Rate | 0.001 |  |  |  |  |  |
| epsilon | 0.01 |  |  |  |  |  |
| Batch Size | 128 |  |  |  |  |  |
| # of Epochs | max. 500 |  |  |  |  |  |
| omega (used for different group lasso weights for different GPs) | 0 for PanglaoDB GPs | 1 for Reactome GPs |  |  |  |  |
| alpha_kl (KL term weight) | 0.1 |  |  |  |  |  |
| alpha_epoch_anneal (epochs for alpha_kl annealing) | 100 (reference) | - |  |  |  |  |

**Supplementary Table 11.** expiMap detailed architecture for the PBMC IFN-*β* deleted genes recovery experiment (**supplementary figure 10**).

| Name | Operation | NoF/Kernel Dim. | Dropout | LN | Activation | Input |
| --- | --- | --- | --- | --- | --- | --- |
| Inputs |  |  |  |  |  |  |
| data | - | #Genes | × | × | - | - |
| conditions | - | #Conditions | × | × | - | - |
| Encoder |  |  |  |  |  |  |
| Layer_1 | FC | 256 | 0.05 | ✓ | ReLU | [data, study labels] |
| Layer_2 | FC | 256 | 0.05 | ✓ | ReLU | Layer_1 |
| Layer_3 | FC | 256 | 0.05 | ✓ | ReLU | Layer_2 |
| mean | FC | #Gene Programs | × | × | Linear | Layer_3 |
| var | FC | #Gene Programs | × | × | Linear | Layer_3 |
| latent | Multivariate Normal | #Gene Programs | × | × | - | [mean, var] |
| Decoder |  |  |  |  |  |  |
| Layer_1 | FC | #Genes | × | × | softmax | [latent, study labels] |
| predicted count mean | Multiplication | #Genes | × | × | - | [Layer_1, library scale] |
| Hyperparameters |  |  |  |  |  |  |
| Loss | NB |  |  |  |  |  |
| Optimizer | Adam |  |  |  |  |  |
| Learning Rate | 0.001 |  |  |  |  |  |
| epsilon | 0.01 |  |  |  |  |  |
| Batch Size | 128 |  |  |  |  |  |
| # of Epochs | max. 200 |  |  |  |  |  |
| alpha (group lasso weight) | 0.7 |  |  |  |  |  |
| alpha_kl (KL term weight) | 0.06 |  |  |  |  |  |
| alpha_epoch_anneal (epochs for alpha_kl annealing) | 100 |  |  |  |  |  |
| alpha_l1 (soft mask weight) | 0.4, 0.3, 0.2, 0.1, 0.06 |  |  |  |  |  |

**Supplementary Table 12.** expiMap detailed architecture for the integration experiment (**figure 3c**, **supplementary figure 7**).

| Name | Operation | NoF/Kernel Dim. | Dropout | LN | Activation | Input |
| --- | --- | --- | --- | --- | --- | --- |
| Inputs |  |  |  |  |  |  |
| data | - | #Genes | × | × | - | - |
| conditions | - | #Conditions | × | × | - | - |
| Encoder |  |  |  |  |  |  |
| Layer_1 | FC | 256 | 0.05 | ✓ | ReLU | [data, study labels] |
| Layer_2 | FC | 256 | 0.05 | ✓ | ReLU | Layer_1 |
| Layer_3 | FC | 256 | 0.05 | ✓ | ReLU | Layer_2 |
| mean | FC | #Gene Programs | × | × | Linear | Layer_3 |
| var | FC | #Gene Programs | × | × | Linear | Layer_3 |
| latent | Multivariate Normal | #Gene Programs | × | × | - | [mean, var] |
| Decoder |  |  |  |  |  |  |
| Layer_1 | FC | #Genes | × | × | softmax | [latent, study labels] |
| predicted count mean | Multiplication | #Genes | × | × | - | [Layer_1, library scale] |
| Hyperparameters |  |  |  |  |  |  |
| Loss | NB |  |  |  |  |  |
| Optimizer | Adam |  |  |  |  |  |
| Learning Rate | 0.001 |  |  |  |  |  |
| epsilon | 0.01 |  |  |  |  |  |
| Batch Size | 128 |  |  |  |  |  |
| # of Epochs | max. 500 |  |  |  |  |  |
| alpha (group lasso weight) | 0.7 |  |  |  |  |  |
| alpha_kl (KL term weight) | 0.1 |  |  |  |  |  |
| alpha_epoch_anneal (epochs for alpha_kl annealing) | 100 |  |  |  |  |  |

**Supplementary Table 13.** Non-amortized scVI detailed architecture for the integration experiment (**figure 3c**, **supplementary figure 7**).

| Name | Operation | NoF/Kernel Dim. | Dropout | BN | Activation | Input |
| --- | --- | --- | --- | --- | --- | --- |
| Inputs |  |  |  |  |  |  |
| data | - | #Genes | × | × | - | - |
| conditions | - | #Conditions | × | × | - | - |
| Encoder |  |  |  |  |  |  |
| mean | Parameter | #Cells × 10 | × | × | - | - |
| var | Parameter | #Cells × 10 | × | × | - | - |
| latent | Multivariate Normal | 10 | × | × | - | [mean, var] |
| Decoder |  |  |  |  |  |  |
| Layer_1 | FC | 128 | × | ✓ | ReLU | [latent, study labels] |
| px_r_decoder | FC | #Genes | × | × | Linear | Layer_1 |
| px_dropout_decoder | FC | #Genes | × | × | Linear | Layer_1 |
| px_scale_decoder | FC | #Genes | × | × | softmax | Layer_1 |
| predicted count mean | Multiplication | #Genes | × | × | - | [px_scale_decoder, library scale] |
| Hyperparameters |  |  |  |  |  |  |
| Loss | ZINB |  |  |  |  |  |
| Optimizer | Adam |  |  |  |  |  |
| Learning Rate | 0.05 |  |  |  |  |  |
| epsilon | 0.01 |  |  |  |  |  |
| Batch Size | 256 |  |  |  |  |  |
| # of Epochs | 600 |  |  |  |  |  |
| Weight Decay | 1e-6 |  |  |  |  |  |
| mean init | PCA |  |  |  |  |  |

**Supplementary Table 14.**
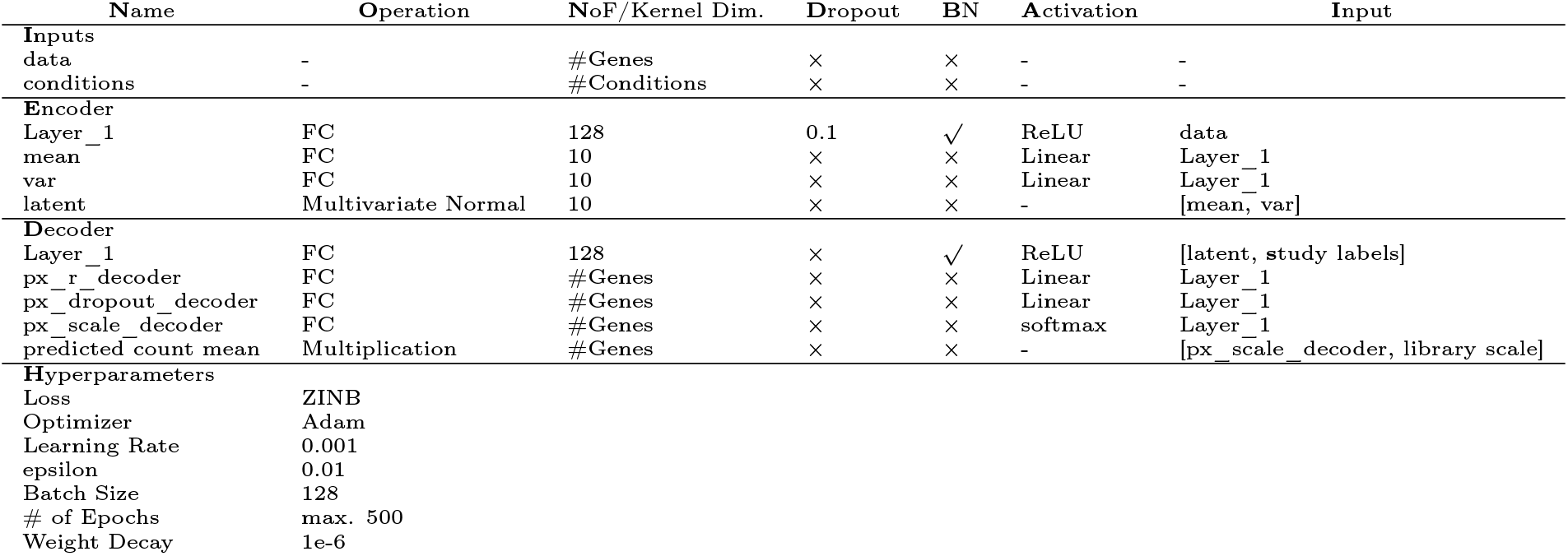
scVI detailed architecture for the integration experiment (**figure 3c**, **supplementary figure 7**).

| Name | Operation | NoF/Kernel Dim. | Dropout | BN | Activation | Input |
| --- | --- | --- | --- | --- | --- | --- |
| **Inputs** |  |  |  |  |  |  |
| data | - | #Genes | × | × | - | - |
| conditions | - | #Conditions | × | × | - | - |
| **Encoder** |  |  |  |  |  |  |
| Layer_1 | FC | 128 | 0.1 | ✓ | ReLU | data |
| mean | FC | 10 | × | × | Linear | Layer_1 |
| var | FC | 10 | × | × | Linear | Layer_1 |
| latent | Multivariate Normal | 10 | × | × | - | [mean, var] |
| **Decoder** |  |  |  |  |  |  |
| Layer_1 | FC | 128 | × | ✓ | ReLU | [latent, study labels] |
| px_r_decoder | FC | #Genes | × | × | Linear | Layer_1 |
| px_dropout_decoder | FC | #Genes | × | × | Linear | Layer_1 |
| px_scale_decoder | FC | #Genes | × | × | softmax | Layer_1 |
| predicted count mean | Multiplication | #Genes | × | × | - | [px_scale_decoder, library scale] |
| **Hyperparameters** |  |  |  |  |  |  |
| Loss | ZINB |  |  |  |  |  |
| Optimizer | Adam |  |  |  |  |  |
| Learning Rate | 0.001 |  |  |  |  |  |
| epsilon | 0.01 |  |  |  |  |  |
| Batch Size | 128 |  |  |  |  |  |
| # of Epochs | max. 500 |  |  |  |  |  |
| Weight Decay | 1e-6 |  |  |  |  |  |

**Supplementary Table 15.** Linear scVI (LDVAE) detailed architecture for the integration experiment (**supplementary figure 7**).

| Name | Operation | NoF/Kernel Dim. | Dropout | BN | Activation | Input |
| --- | --- | --- | --- | --- | --- | --- |
| **Inputs** |  |  |  |  |  |  |
| data | - | #Genes | × | × | - | - |
| conditions | - | #Conditions | × | × | - | - |
| **Encoder** |  |  |  |  |  |  |
| Layer_1 | FC | 128 | 0.1 | ✓ | ReLU | data |
| mean | FC | 10 | × | × | Linear | Layer_1 |
| var | FC | 10 | × | × | Linear | Layer_1 |
| latent | Multivariate Normal | 10 | × | × | - | [mean, var] |
| **Decoder** |  |  |  |  |  |  |
| px_dropout_decoder | FC | #Genes | × | ✓ | Linear | [latent, study labels] |
| factor_regressor | FC | #Genes | × | ✓ | softmax | [latent, study labels] |
| predicted count mean | Multiplication | #Genes | × | × | - | [factor_regressor, library scale] |
| **Hyperparameters** |  |  |  |  |  |  |
| Loss | NB |  |  |  |  |  |
| Optimizer | Adam |  |  |  |  |  |
| Learning Rate | 0.001 |  |  |  |  |  |
| epsilon | 0.01 |  |  |  |  |  |
| Batch Size | 128 |  |  |  |  |  |
| # of Epochs | max. 500 |  |  |  |  |  |
| Weight Decay | 1e-6 |  |  |  |  |  |

**Supplementary Table 16.**
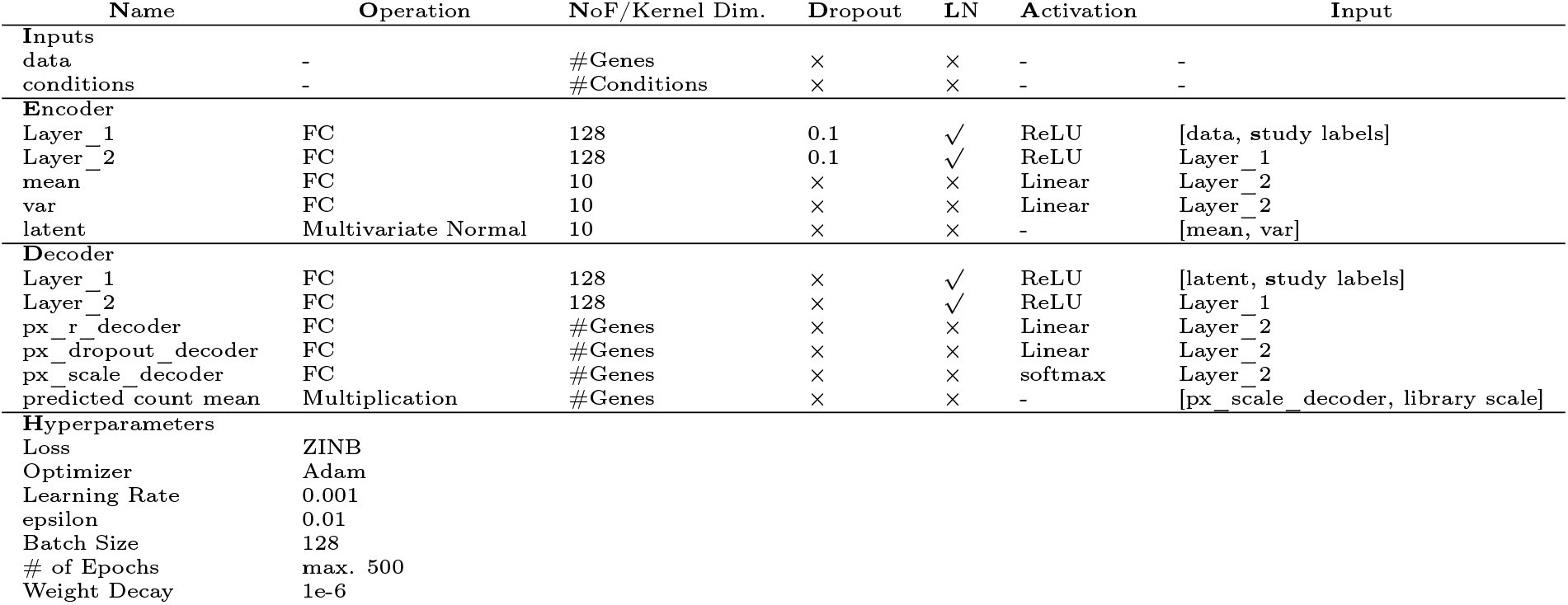
scVI detailed architecture for the query to reference projection (**figure 3a, b** and **supplementary figure 5**, **supplementary figure 13b**).

## Notes

https://github.com/theislab/expiMap_reproducibility

### Competing Interest Statement

Fabian J. Theis consults for Immunai Inc., Singularity Bio B.V., CytoReason Ltd, and Omniscope Ltd, and has ownership interest in Dermagnostix GmbH and Cellarity.

